# A visual atlas of genes’ tissue-specific pathological roles

**DOI:** 10.1101/2022.01.08.475476

**Authors:** Priyadarshini Rai, Atishay Jain, Neha Jha, Divya Sharma, Shivani Kumar, Abhijit Raj, Apoorva Gupta, Sarita Poonia, Smriti Chawla, Angshul Majumdar, Tanmoy Chakraborty, Gaurav Ahuja, Debarka Sengupta

## Abstract

Dysregulation of a gene’s function, either due to mutations or impairments in regulatory networks, often triggers pathological states in the affected tissue. Comprehensive mapping of these apparent gene-pathology relationships is an ever daunting task, primarily due to genetic pleiotropy and lack of suitable computational approaches. With the advent of high throughput genomics platforms and community scale initiatives such as the Human Cell Landscape (HCL) project [1], researchers have been able to create gene expression portraits of healthy tissues resolved at the level of single cells. However, a similar wealth of knowledge is currently not at our finger-tip when it comes to diseases. This is because the genetic manifestation of a disease is often quite heterogeneous and is confounded by several clinical and demographic covariates. To circumvent this, we mined ~18 million PubMed abstracts published till May 2019 and selected ~6.1 million of them that describe the pathological role of genes in different diseases. Further, we employed a word embedding technique from the domain of Natural Language Processing (NLP) to learn vector representation of entities such as genes, diseases, tissues, etc., in a way such that their relationship is preserved in a vector space. Notably, Pathomap, by the virtue of its underpinning theory, also learns transitive relationships. Pathomap provided a vector representation of words indicating a possible association between *DNMT3A*/*BCOR* with *CYLD* cutaneous syndrome (CCS). The first manuscript reporting this finding was not part of our training data.

**Key points:**

- We mined ~18 million PubMed abstracts to extract latent knowledge pertaining to tissue specific pathological roles of genes.
- We found well-defined gene modules implicated in disease pathogenesis in anatomically proximal tissues.
- We demonstrated an ahead of time discovery of the association between DNMT3A/BCOR with CYLD cutaneous syndrome (CCS), as a knowledge synthesis use-case.

## Background

The number of published biomedical manuscripts has sharply increased since the beginning of the genomic era. Nearly 3000 manuscripts are published daily in peer-reviewed journals [2]. This has resulted in a voluminous corpus of scientific literature, outreaching by far, the limit of human comprehension. Given the complexity of human biological systems, it is nearly impossible to get a qualitative as well as a quantitative view of organ/tissue wide pathological roles of genes by an internet search of the literature. Today, a great deal of community level effort is channeled towards creating a molecular atlas of healthy human organs by charting out the functionally distinct cellular subtypes in tissues of interest. Such effort is scarce in disease biology due to the presence of innumerable covariates and the diversity of the disease states. A vast majority of knowledge-based archiving disease-gene association is painstakingly, manually curated. DisGeNET [3] and Online Mendelian Inheritance in Man (OMIM®) [4] are noteworthy in this regard. NLP based efforts in this regard have mainly concentrated on extraction of disease-gene association from biomedical literature which is an automated version of manual data curation [5]. While these repositories are imperative to take our disease biology investigations forward, none of these are equipped to give a comprehensive atlas-like view of genes’ tissue specific pathological roles.

Here we present Pathomap, a visual atlas of genes’ tissue-specific pathological roles. To achieve this, we took the help of word embedding technologies from the domain of NLP. Word embedding typically employs efficient neural networks to learn word-representations by ingesting millions of documents. Mikolov and colleagues first proposed the very idea of learning a continuous bag of representation of words at Google LLC [6]. The field has since evolved dramatically, and in recent times, transformer based models such as BERT (bidirectional encoder representations from transformers) [7] have emerged as state-of-the-art. In the past years, text mining and language modeling have gained prominence in the fields of biomedical research. Gideon et al. used word embedding techniques to identify associations among different brain regions [8]. Similar approaches have been used in predicting compound-protein interactions [9] and drug discovery [10]. BERT is highly resource intensive. In a recent effort by Lee and colleagues, embeddings have been produced for several billions of words by processing voluminous biomedical literature, using BERT [2]. We hypothesized that such generic word embeddings may not capture the essence of molecular pathology. To circumvent this, we first screened ~6.1 million abstracts that describe an association between certain genes’ dysregulation and diseases from a database of ~18 million biomedical abstracts. Word embeddings were learned exclusively on the basis of these selected abstracts.

In contrast to the generic BioBERT embeddings, the Pathomap word vectors were found to preserve semantic relationships between words in a manner that is consistent with our knowledge of the pathogenesis of diverse diseases. Furthermore, these disease-focused word embeddings allowed us to extract latent knowledge beyond the ambit of the training literature corpus.

## Materials and Methods

### Data Description

We downloaded ~18 million abstracts [11] from PubMed that were published till May 2019. We postulated that abstracts capture the essence of the entire manuscript [28]. Also, due to paywalls, many articles are not accessible for mining.

### Manual annotation of abstracts

We looked for articles in the field of molecular biology and genetics, and thereby manually curated two sets of abstracts - first, describing genes’ direct roles in disease pathogenesis, and second, abstracts that do not cite apparent disease-gene relationships. The former consisted of 1,363 abstracts (**Supplementary File 1**) and the latter consisted of 768 abstracts (**Supplementary File 2**). The abstracts with information about gene-disease relationships were labeled as ‘relevant’ or ‘positive’ data and the remaining abstracts that did not mention gene and corresponding disease relationships were labeled as ‘non-relevant’ or ‘negative’ data. We ensured that the number of positively and negatively annotated abstracts is approximately the same in number. For the annotation of abstracts, we also referred to articles from GWAS (genome-wide association study), COSMIC(Catalogue Of Somatic Mutations In Cancer), and OMIM (Online Mendelian Inheritance in Man) databases.

### Stratifying pathological abstracts from non-pathological abstracts

An abstract stratification strategy was developed to shortlist the abstracts that depict the pathological role of genes. We used positive and negative sets of abstracts as seed data. These abstracts were further divided into train and test sets in a 75:25 split. We used BioSentVec [12] to transform these abstracts into vectors of dimension 700 that are suitable for machine learning. We evaluated the performance of three classifiers, namely, Support Vector Machine (SVM) [13], Extreme Gradient Boosting (XGBoost) [14], and Logistic Regression (LR) [15] on the associated binary classification task. SVM outperformed the other two methods in terms of prediction accuracy (**Supplementary Table S1, Supplementary Table S2, Supplementary Figure S1**). We classified all ~18 million abstracts using the SVM model (https://github.com/Priyadarshini-Rai/Pathomap/tree/main/Pathomap/inst/extdata). As a stringent selection criterion, we qualified an abstract as *patho-abstract* if the SVM model returned prediction probability was greater than 0.7. After classification, we obtained ~6.1 million abstracts as *patho-abstract*.

### Word2vec methodology

The ~6.1 million abstracts obtained after classification were subjected to train the Word2vec skip-gram model. We performed pre-processing by first performing tokenization using python’s nltk’s word tokenizer. After that, we removed punctuations and stop-words from the list of tokens. We found that lower-casing the articles helped reduce the computational overhead and vocabulary size without significantly affecting the performance. We finally used the *patho-abstracts* to train our primary Word2vec model. Phrases provided by ‘gensim’ in python were used to handle bigrams. Window size and vector length were considered to be 5, and 300 respectively. Word embeddings trained on *patho-abstracts* are shared on Zenodo (https://zenodo.org/record/5237312). A tissue specific *Patho-score* is calculated simply as the cosine distance between the vectors associated with a tissue name and a certain gene symbol. **Figure 1A** outlines the method workflow.

**Figure 1:**
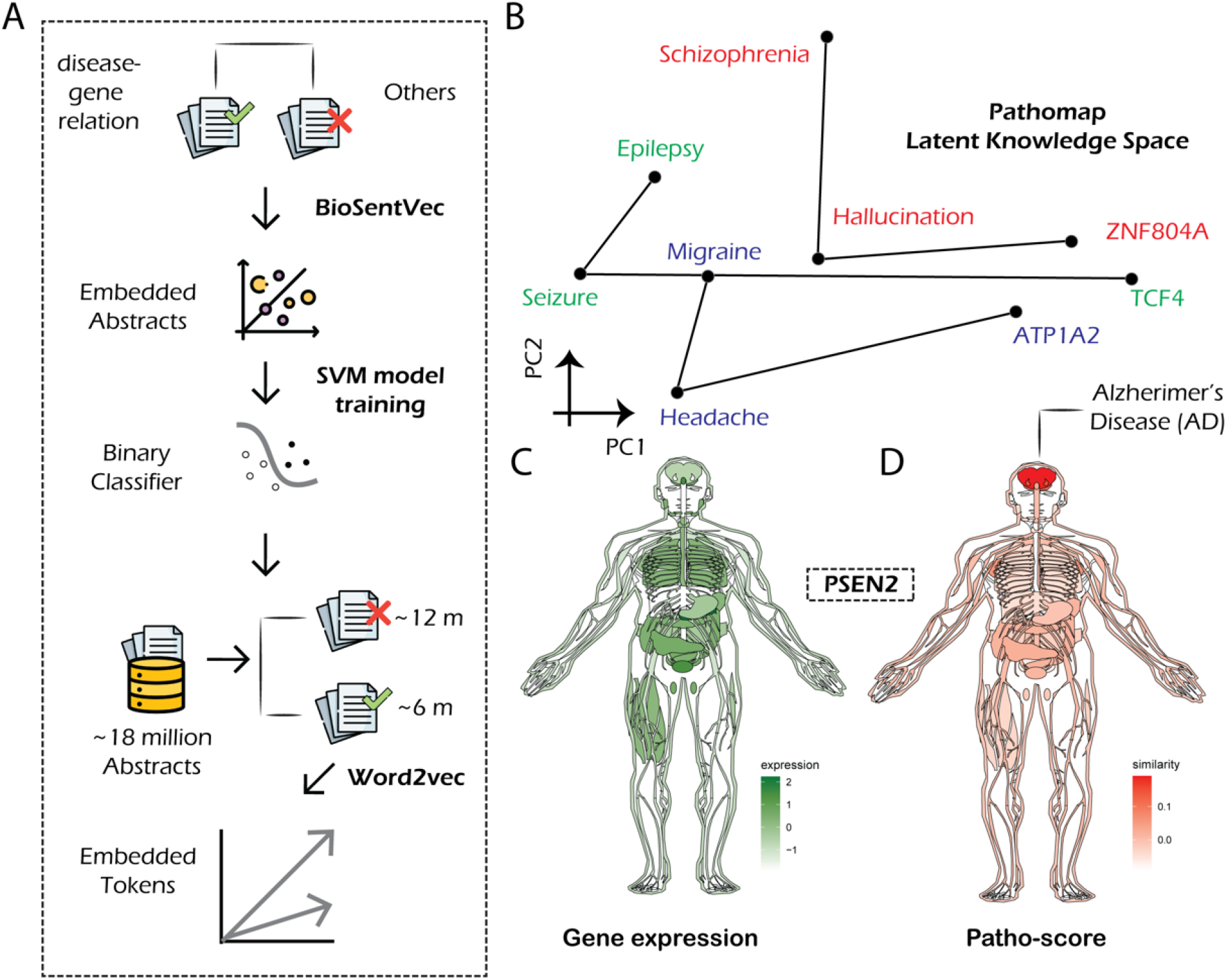
(A) Steps involved in representation learning from abstracts describing disease-gene associations. (B) Congruence of disease-symptom-causative gene triplets (schizophrenia-hallucination-ZNF804A; epilepsy-seizure-TCF4; migraine-headache-ATP1A2) in the word-vector space. (C) Expression and pathological role of *PSEN2* are shown as an example, on the human body template. While tissue wide expression levels of the gene are largely uniform, *Patho-score* shows remarkably higher intensity in the brain. Notably, mutations in *PSEN2* are the cause of familial Alzheimer’s disease.

### Pathomap for visualizing the impact of a gene on different tissues

For wide dissemination and ease of use, we created a Shiny application (https://the-ahuja-lab.shinyapps.io/Pathomap), which is freely available to users. An R package is also compiled to give users programmatic access to the embeddings. Both these mediums allow users to visualize a gene’s tissue specific pathogenic roles and compare them with tissue-wide expression levels, obtained from GTEx gene expression archives [16]. We will update the Shiny application and R package yearly to incorporate new discoveries. Of note, we used the gganatogram R package to visualize *Patho-scores* on a human body template [17]. For visualizing gene expression levels, we applied median normalization and log transformation to the GTEx expression matrix.

## Insights and Applications

### Semantic similarity among word-relationships

To evaluate the meaningfulness of the word vectors, we constructed a case-study to visualize the semantic similarity between disease-symptom-gene relationships. We considered three such triplets — schizophrenia-hallucination-ZNF804A [18]; epilepsy-seizure-TCF4 [19]; migraine-headache-ATP1A2 [20]. We visualized vectors associated with these words using Principal Component Analysis (PCA) (**Figure 1B**). The relative positioning of the words across the triplets exhibits consistent directionalities, such that there exist consistent vector operations between words that represent concepts such as ‘symptom of’ and ‘causative gene for’. Vectors associated with about 2 million words, trained on *patho-abstracts* are made available on Zenodo (https://doi.org/10.5281/zenodo.5237312). These can be used as an exploratory tool to visualize, discover, and confirm more such relationships.

### Disease associated genes are shared among anatomically proximal tissues

Diseases are best understood in terms of the dysregulation of genes and other regulatory components. It’s not convenient to explore if genes share tissue specific disease associations. To validate this, we plotted a heatmap depicting 1000 genes with the highest standard deviation of *Patho-score* across 34 main tissue types (**Figure 2**). We see the emergence of neat gene modules with reproducible intensity patterns of *Patho-scores* across diverse tissues. For example, all important brain regions (hypothalamus, cerebellum, frontal cortex, amygdala, and hippocampus) share the disease causing genes. Similarly, skeletal muscle and adipose tissue are found to share the disease associated genes. Notably, metabolic interrelationships and cross-talk of signals derived from both skeletal muscle and adipose tissue have been long investigated [21]. The prostate and the breast are two organs with several similarities: lobular glands, gonadal steroid sensitivity, and diseases treated by some comparable therapeutic approaches, primarily in the context of cancer. In fact, pathological evaluation of these two organs is similar in nature [22]. Both these organs shared similar patterns associated with *Patho-scores*. We failed to see such meaningful structures when we attempted to reproduce this heatmap using pre-trained BioBERT embeddings (**Supplementary Figure S2**). This highlights the power of context specific learning of word-vectors.

**Figure 2:**
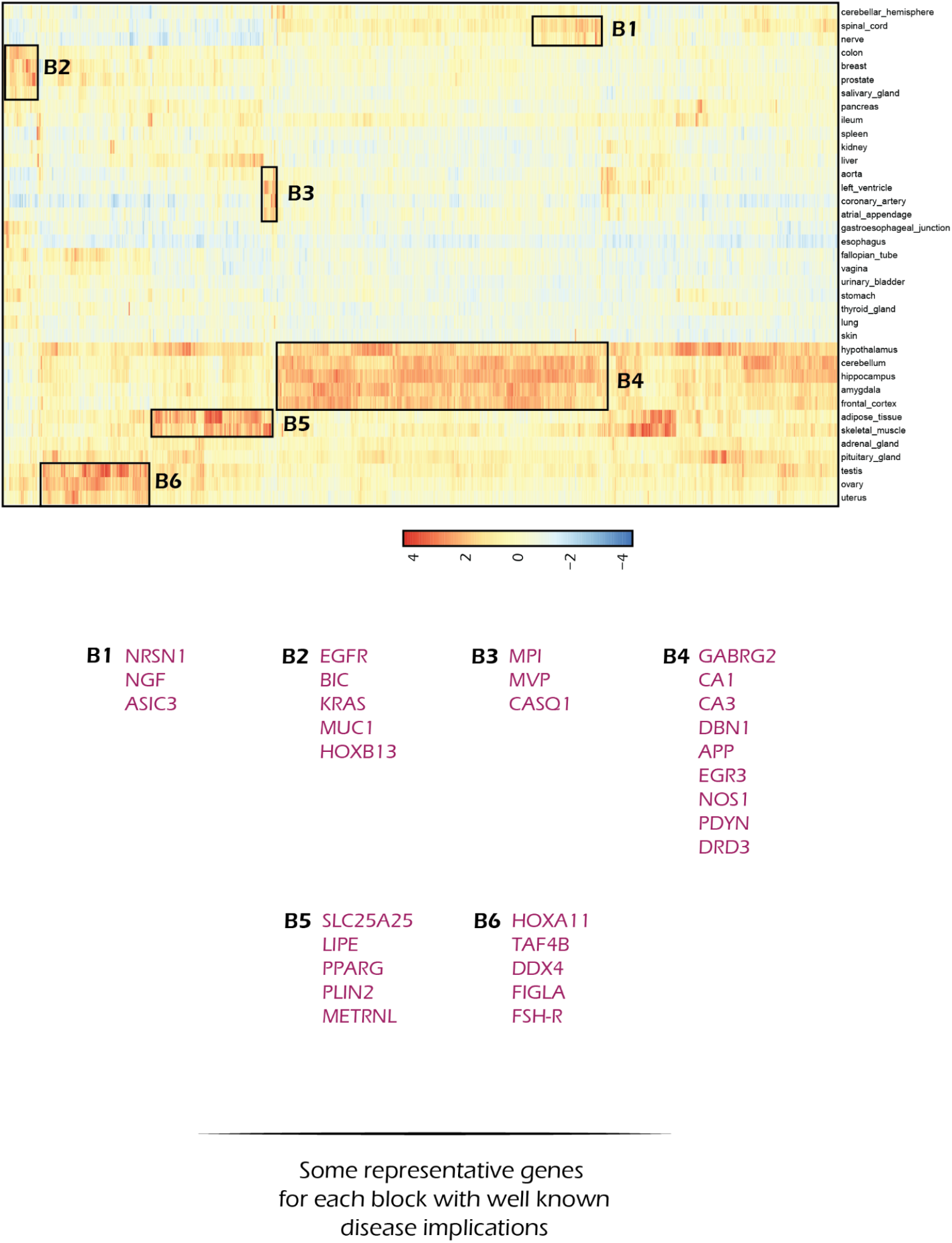
Heatmap depicting *Patho-scores* indicating tissue specific pathological involvement of genes. Each column of the heatmap represents one of the 1000 genes having the highest standard deviation of *Patho-scores* across tissues. For each block (manually boxed on the basis of coherent color intensities) some representative genes are shown with known pathological implications in concerned tissues.

### Disease linked genes are uniformly distributed across the human genome

We speculated that there could be genomic hotspots of disease linked genes. To test this, we divided the length of each of the human chromosomes into contiguous chunks spanning 1 million base pairs and counted the total number of genes located in each of these DNA stretches. Further, we counted the number of times gene(s) located in each of these stretches appeared in the *patho-abstracts*. We found these values to be highly correlated across the 1 million base-pair long DNA stretches (**Supplementary Figure S3**) with a Spearman’s correlation coefficient of 0.7.

### Visual atlas of disease linked genes

Since the beginning of the genomic era, several studies have been conducted to create molecular portraits of healthy tissues using transcriptomic, proteomic, and epigenomic assays. Single cell transcriptomics has fueled such efforts by providing unprecedented resolution into phenotypic heterogeneity in cells. With the advent of a high throughput single-cell transcriptomics platform, the past few years have seen a sharp spike in single cell atlasing effort, disentangling versatile molecular architecture of healthy tissues. It is difficult to undertake such studies for diseased tissues due to the non-deterministic nature of genetic aberrations that underpin their associated pathogenesis. Pathomap is the first unbiased, metainformation platform to map genes’ pathogenic role at the tissue level. We developed an R package as well as a Shiny web server to facilitate gene queries. For example, **Figure 1C** and **Figure 1D** depict *PSEN2* expression and *Patho-score* respectively across human tissues. Mutations in the *PSEN2* gene are known to have a causal role in Alzheimer’s disease [23]. When it comes to gene expression, *PSEN2* is minimally variable across tissues. Several, well-known disease linked genes were tested to ensure the integrity of the platform (**Supplementary Figures S4-19**)

### Word analogy and knowledge synthesis

Word embeddings can be used to find domain specific analogies. We tested various well-known relationships. For instance, the most similar gene to diabetes associated with *TCF7L2* [24] was found to be *CDKAL1* with a cosine similarity of 0.69. Notably, *CDKAL1* is also implicated in type 2 diabetes [25]. Second, we asked, “diabetes is to the pancreas, as asthma is to?” We obtained trachea, bronchi, lung as top hits with a cosine similarity of 0.56, 0.56, and 0.52 respectively. To the question “diabetes is to diuresis as asthma is to?”, we got the answer as bronchoconstriction, cough, bronchospasm with a cosine similarity of 0.58, 0.54, and 0.53 respectively. We also tried to predict the type of cancer based on the gene — “*BRCA* is to breast cancer as *KRAS* is to which cancer?”, where the most similar response given by the embedding is colorectal cancer having a cosine similarity score of 0.62 [26]. *TCF7L2* variants have been associated with type 2 diabetes in multiple ethnic groups [24]. “Diabetes is caused by aberrations in *TCF7L2* as schizophrenia is caused by which gene?” Among all genes, *ZNF804A* was returned as the top gene [18]. Such analogies can be represented through algebraic equations involving concerned word-vectors as follows.

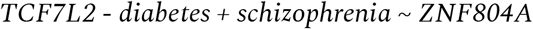

Word embeddings are capable of deriving latent knowledge. The latest publication in our abstract repository was dated May 2019. A paper by Davies et al., published on Oct 17, 2019, reported molecular profiling data of the genomic landscape (42 benign and malignant tumors across 13 individuals from four multigenerational families) and discovered recurrent mutations in epigenetic modifiers *DNMT3A* and *BCOR* in 29% of benign tumors [27]. To the best of our knowledge, this was the first report linking these genes with *CYLD* cutaneous syndrome (CCS). We obtained high cosine similarity between these genes and *CYLD*: (‘CYLD’,’DNMT3A’) = 0.30 (‘CYLD’,’BCOR’) = 0.37.

### Benchmarking of Pathomap’s automated approach against manual curation

DisGeNET features a large number of disease-gene relationships, which can be considered as ground truth data [3]. We found 9727 such relationships for which disease and gene names were present among the tokens for which word embeddings were produced. These are considered positive samples. An equal number of randomly disease-gene pairs were populated and associated *Patho-scores* were obtained. These *Patho-scores* are considered as negative samples. We hypothesized that if Pathomap’s automated approach is viable, we should typically expect clearly higher *Patho-scores* for experimentally validated disease-gene relationships, logged by DisGeNET. We compared the distribution of *Patho-scores* associated with positive and negative samples (**Figure 3A**). The *Patho-scores* associated with the positive samples showed significant right skew in distribution as compared to negative samples, depicting high fidelity of the *Patho-scores* above a cut-off of 0.3. We compared the distributions using the Kruskal-Wallis test (*p-value* < 2.2e-16). Pathomap has about half of the DisGeNET annotated diseases in the database **(Figure 3B)**. Further, we downloaded the list of human genes from NCBI and gene-disease associations from the DisGeNET database. The human genes list downloaded from NCBI has ~25K genes. We found ~15K of these genes in *Pathomap*, whereas ~9K genes in the DisGeNET database. ~8.8K genes were common in both databases.

**Figure 3:**
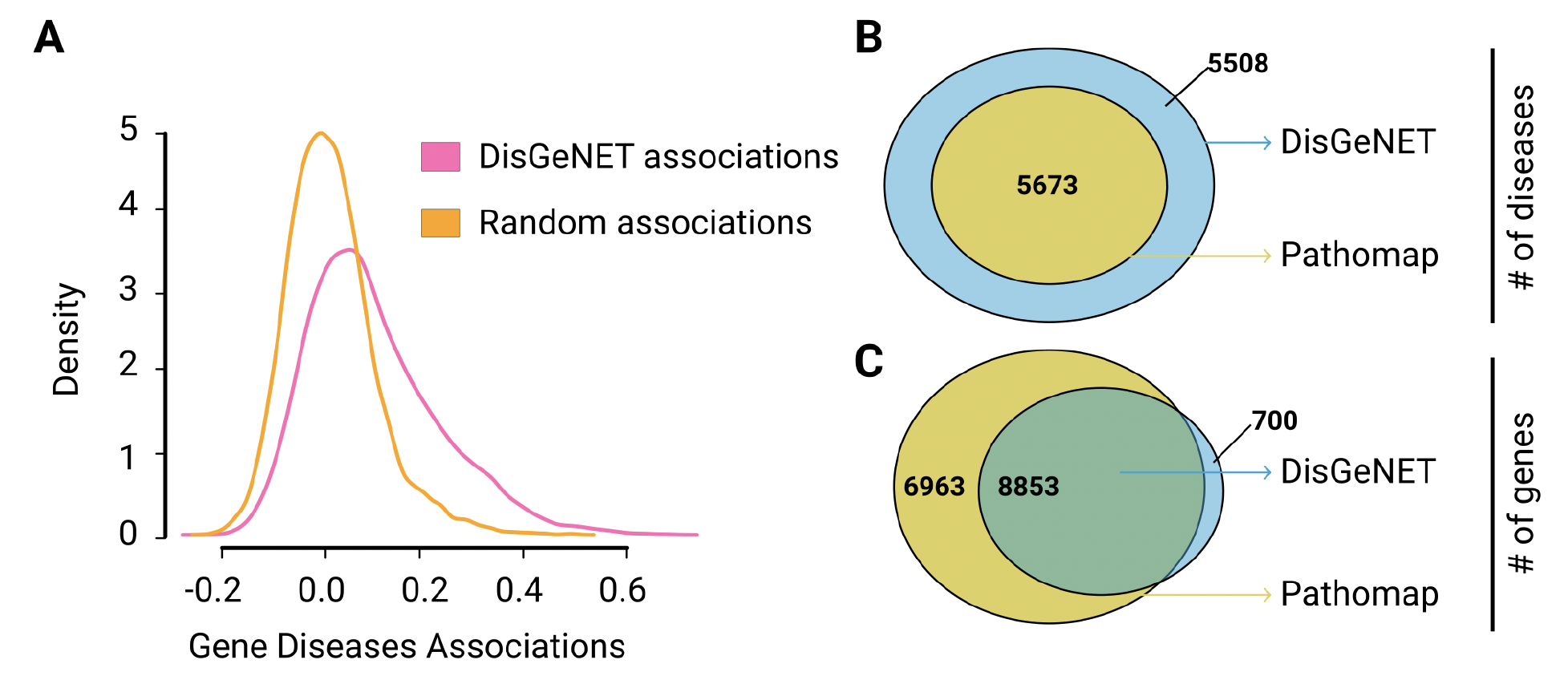
(A) The plot depicts the density distribution of the *Patho-score* of diffierent gene-disease pairs from DisGeNET and compares it with the *Patho-score* distribution of random gene-disease pairs. The density distribution of random gene-disease pairs follows normal distribution whereas the gene-disease pairs from the DisGeNET database are right-skewed depicting the meaningful association between a gene and the corresponding disease. (B) The above figure represents that Pathomap covers more genes that are listed in the NCBI homo sapiens genes list in comparison to DisGeNET. (C) The gene-disease associations from the DisGeNET database has 11,181 disease names out of which 5673 disease names were also present in the Pathomap word embeddings.

## Discussion and Conclusion

Pathomap provides a strategy to obtain an unbiased continuous representation of disease causing genes and their tissue specificity. This can accelerate targeted investigations into diverse diseases. *Patho-scores* and associated color intensities indicate the extent of a gene’s involvement in causing disease in a particular tissue/organ. Pathomap does not provide specifics of the gene’s involvement in a particular disease. For that one can look into PubMed. It is well known that gene expression matrices are low-rank. Our analysis of the top 1000 tissue wide differentially disease-implicated genes suggests that their pathological roles are reproducibly patterned (**Figure 2**). This is a high-level, empirical view and by design, trustworthy.

A major caveat of the current study is that we did not perform species wise segregation of the articles. As such, the current scores do not discriminate between mice, humans, or other primates. We believe this won’t introduce any nuisance factors since animal models are typically used to mimic human diseases only. Pathomap can be conveniently used for deciding genes for diagnostic gene-panels. We demonstrated Pathomap’s ability to synthesize previously unknown associations between diseases and genes scopes. Pathomap may also serve as an orthogonal approach for Gene Ontologies for the narrow and precise investigation of outputs from high-throughput experiments such as RNA-Seq or MicroArray. It can also assist in prioritizing the gene selection for the large-scale loss of function studies.

We compared Pathomap’s automated literature mining approach to the high fidelity manual curation approach taken by DisGeNET. We found *patho-scores* to be extremely reliable beyond a moderately conservative cutoff. While human cell atlasing is a reality, a pathological atlas might take significant effort and resources, which we predict to not arrive anytime soon. Till the time this vision becomes a reality, we trust that Pathomap will assist the community in narrowing the investigation scopes.

### Availability

Pathomap is freely available as both a web server and an R software package. Web Server: https://the-ahuja-lab.shinyapps.io/Pathomap/; R package: https://github.com/Priyadarshini-Rai/Pathomap/blob/main/Pathomap/README.md.

## Supporting information

relevant or postive abstracts

non-relevant or negative abstracts

## Acknowledgment

DS conceived the study. PR wrote the code and performed all analyses with assistance from AJ, SK, AR, and AG. NJ, DSH (Divya Sharma), SC, SP performed manual annotations of abstracts. NJ and DSH performed analysis on word analogies and knowledge synthesis. TC and GA supervised NLP related developments, biological interpretation alongside DS. AM assisted in improving the machine learning workflow.

