## supplementary information for "A visual atlas of genes’ tissue-specific pathological roles"

**Supplementary Table S1:** The below table depicts the performance of different classification algorithms achieved while separating the abstracts featuring disease-gene association and ones that are generic in nature. Finally, a support vector classifier (SVC) was used for classification.

| **Model** | **Accuracy** | **Precision** | **Recall** | **F1 Score** | **Cohen's Kappa** |
|---|---|---|---|---|---|
| **Support Vector Classifier (SVC)** | **0.958** | **0.973** | **0.961** | **0.967** | **0.910** |
| **Extreme Gradient Boosting (XGBoost)** | 0.953 | 0.956 | 0.970 | 0.963 | 0.897 |
| **Logistic Regression (LR)** | 0.949 | 0.961 | 0.959 | 0.960 | 0.889 |

**Supplementary Table S2:** The below table represents extensive hyperparameter tuning for the SVM classifier and reports the results for the different hyperparameter combinations. The regularization parameter C is not tested for higher values because we wanted a larger margin between the positive and negative samples so as to distinguish the two classes clearly.

| **Hyperparameters** | | **Accuracy** | **Precision** | **Recall** | **F1-score** | **Cohen Kappa** |
|---|---|---|---|---|---|---|
| **Kernel** | **C (Regularization parameter)** | | | | | |
| polynomial (degree: 1) | 0.1 | 0.9618 | 0.9810 | 0.9593 | 0.9699 | 0.91749 |
| | 0.5 | 0.9630 | 0.9810 | 0.9611 | 0.9708 | 0.92030 |
| | 1.0 | **0.9662** | 0.9820 | 0.9649 | 0.9733 | 0.92702 |
| polynomial (degree: 2) | 0.1 | 0.9355 | 0.9687 | 0.9642 | 0.9663 | 0.90592 |
| | 0.5 | 0.9568 | 0.9687 | 0.9642 | 0.9663 | 0.9059 |
| | 1.0 | 0.9624 | 0.9716 | **0.9700** | 0.9707 | 0.91821 |
| polynomial (degree: 3) | 0.1 | 0.9549 | **0.9829** | 0.9467 | 0.9643 | 0.90294 |
| | 0.5 | 0.9630 | 0.9801 | 0.9624 | 0.9710 | 0.91985 |
| | 1.0 | **0.9662** | 0.9802 | 0.9672 | **0.9736** | **0.92647** |
| polynomial (degree: 4) | 0.1 | 0.9505 | 0.9701 | 0.9526 | 0.9612 | 0.89284 |
| | 0.5 | 0.9561 | 0.9715 | 0.9604 | 0.9658 | 0.90460 |
| | 1.0 | 0.9643 | 0.9764 | 0.9683 | 0.9722 | 0.92216 |
| rbf | 0.1 | 0.9493 | 0.9798 | 0.9410 | 0.9596 | 0.89132 |
| | 0.5 | 0.9599 | 0.9802 | 0.9573 | 0.9683 | 0.91352 |
| | 1.0 | 0.9618 | 0.9822 | 0.9582 | 0.9698 | 0.91756 |
| linear | 0.1 | 0.9505 | 0.9721 | 0.9503 | 0.9609 | 0.89313 |
| | 0.5 | 0.9505 | 0.9721 | 0.9503 | 0.9609 | 0.89313 |
| | 1.0 | 0.9505 | 0.9721 | 0.9503 | 0.9609 | 0.89313 |

| | | | | | | |
|---|---|---|---|---|---|---|
| sigmoid | 0.1 | 0.9612 | 0.9810 | 0.9583 | 0.9693 | 0.91622 |
| | 0.5 | 0.9624 | 0.9811 | 0.9601 | 0.9703 | 0.91889 |
| | 1.0 | 0.9643 | 0.9802 | 0.9640 | 0.9719 | 0.92277 |

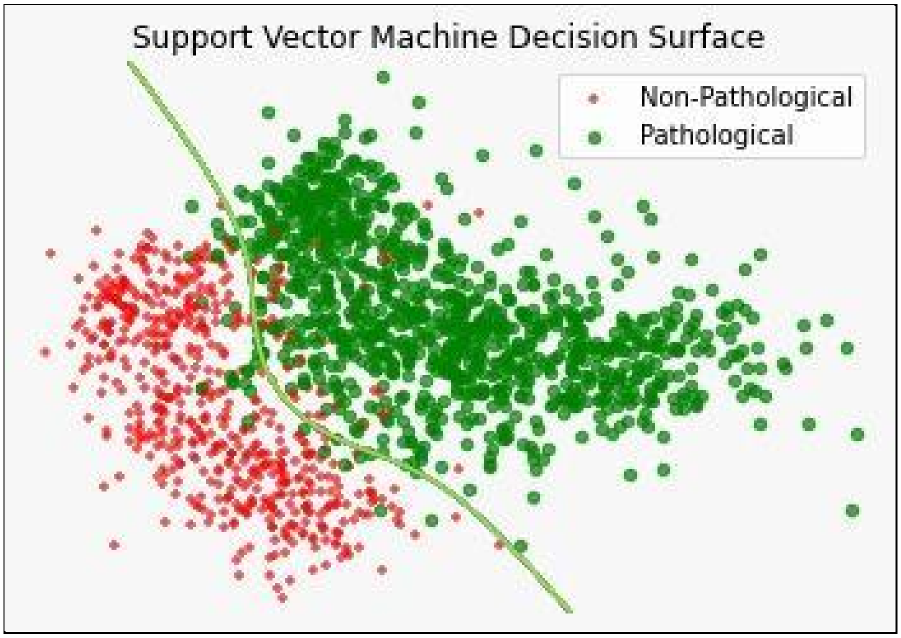


**Supplementary Figure S1.** The above figure represents the support vector classifier's (SVC) decision boundary. The *patho-abstracts* are clearly segregated from ones that are generic.

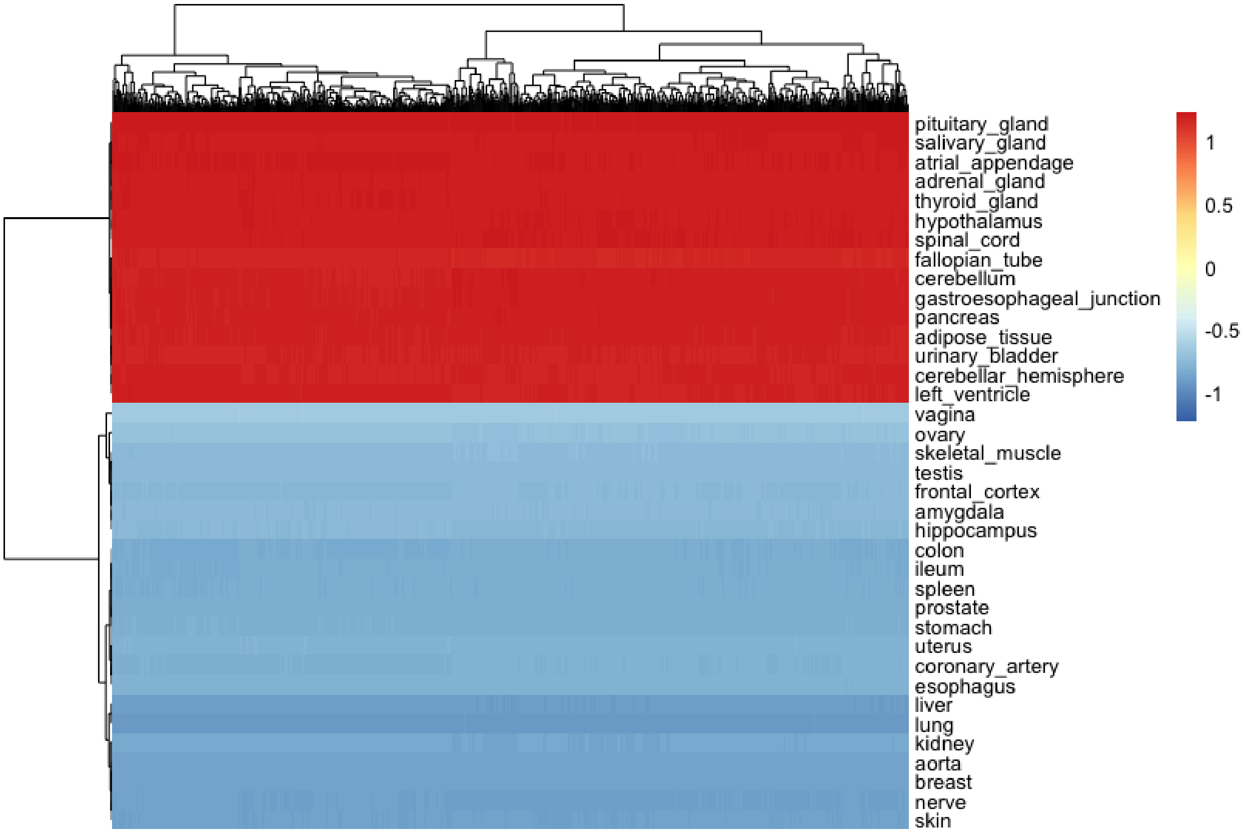


**Supplementary Figure S2.** Heatmap represents cosine similarity between 34 different tissues and top 1000 variable genes (based on standard deviation of the Patho-scores across tissues) using BioBERT embeddings (analogous to Figure 2).

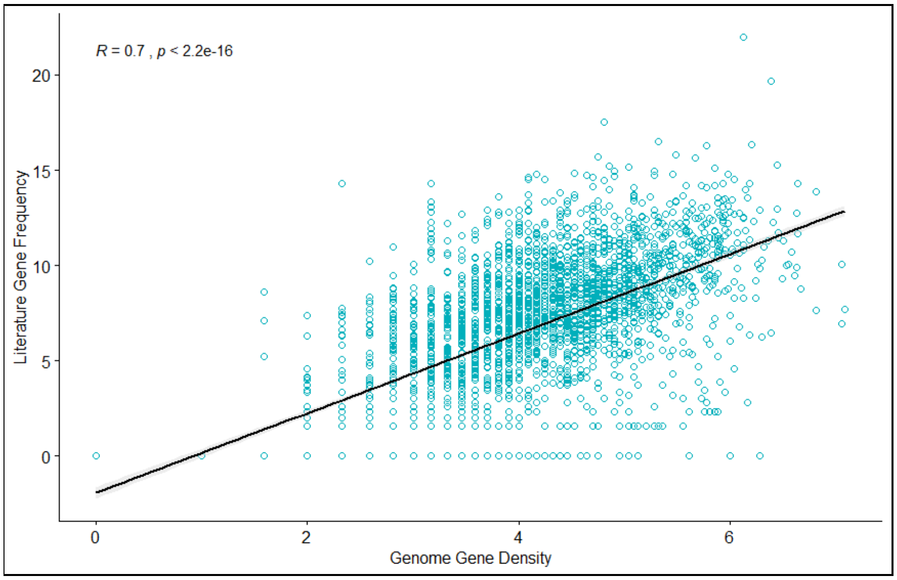


**Supplementary Figure S3.** Correlation between gene numbers across all 1 million bp windows of the human DNA and frequency of their occurrence (genes') in *patho-abstracts*.

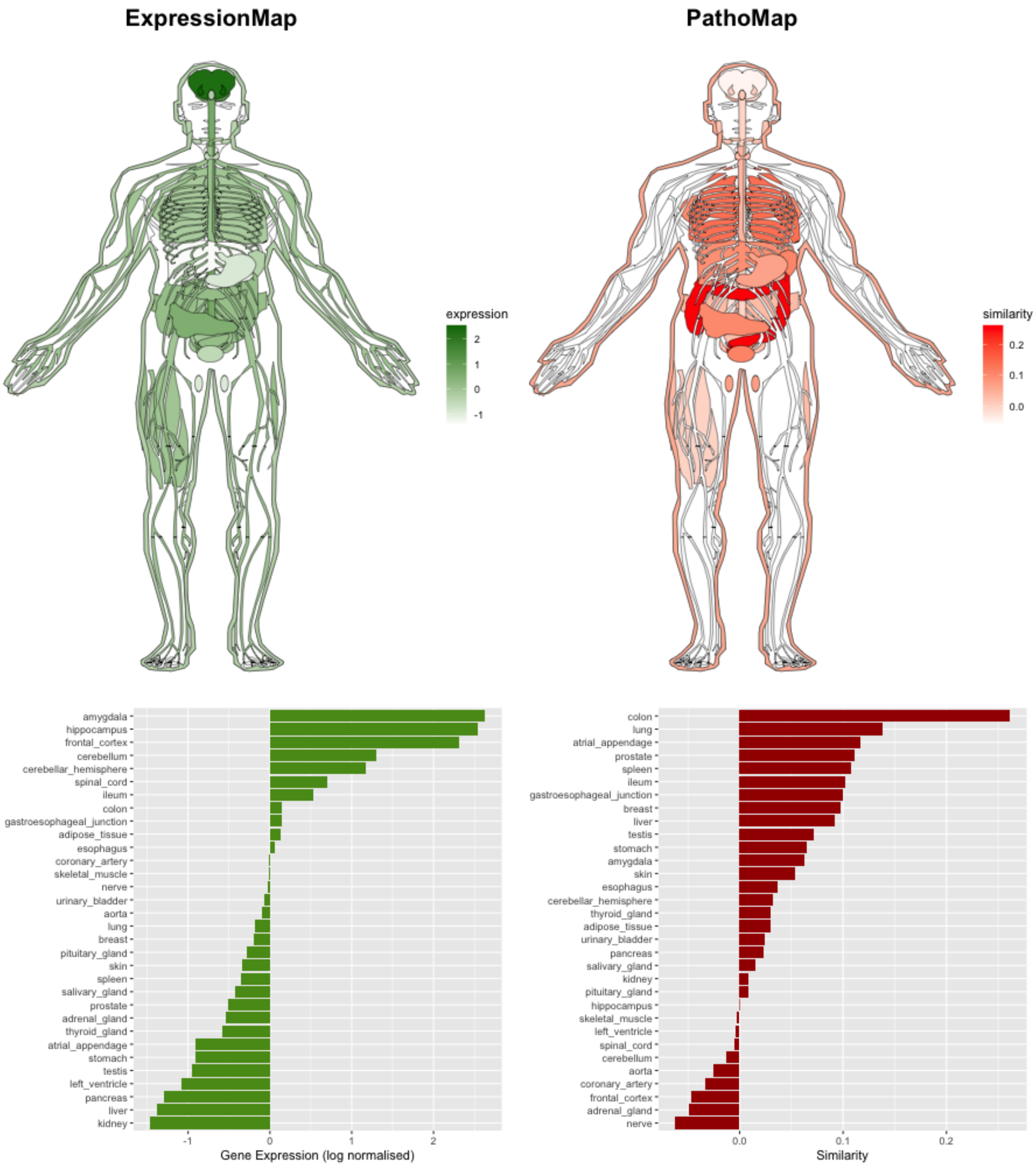


**Supplementary Figure S4.** Distribution of patho-scores and gene expression across organs for APC. The majority of instances of familial adenomatous polyposis (FAP) are caused by germline mutations in the APC gene (Leoz et al. 2015). However, mutations in APC genes are not only limited to FAP but they also play a rate-limiting role in colorectal tumors (Fodde 2002).

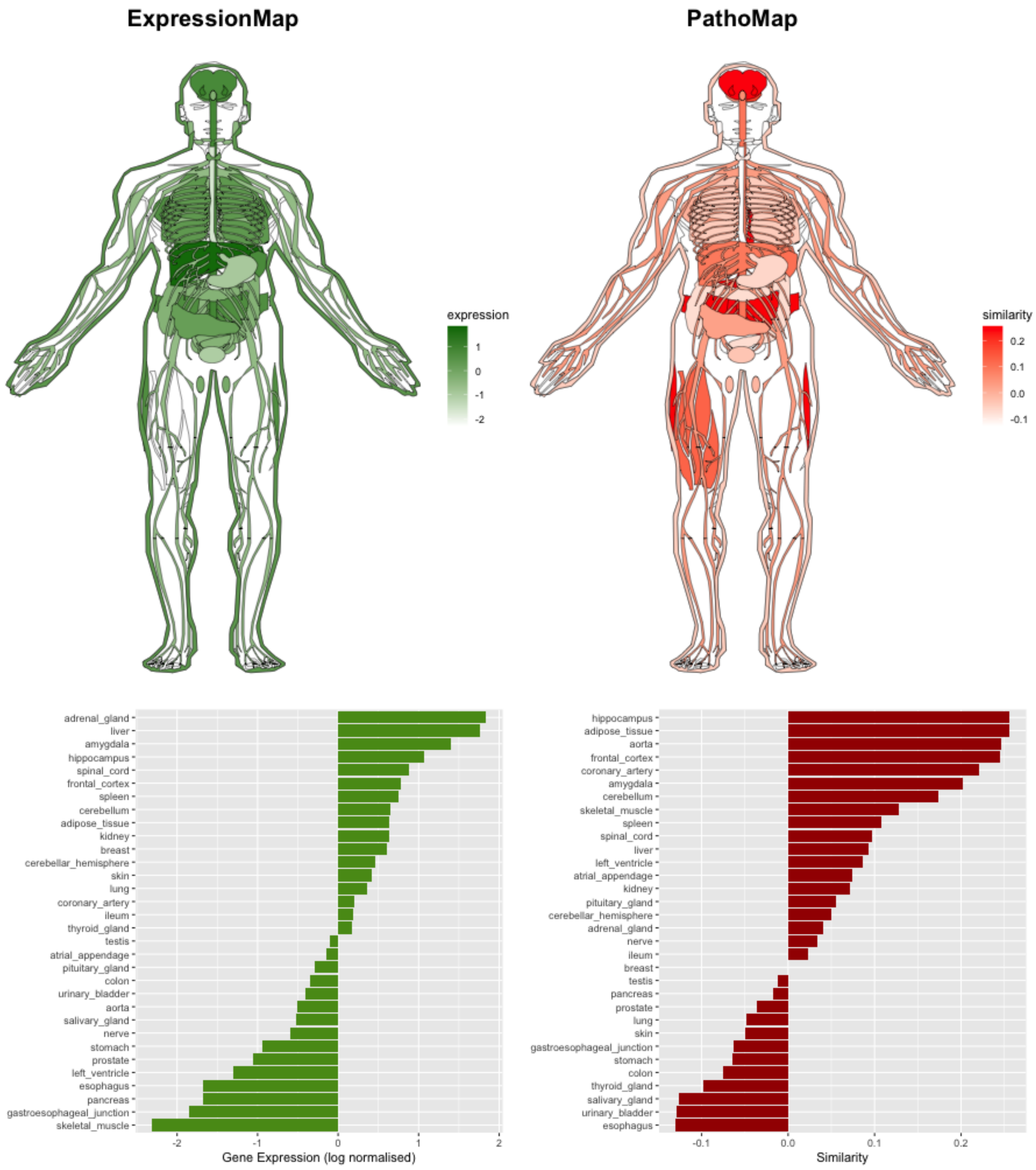


**Supplementary Figure S5.** Distribution of patho-scores and gene expression across organs for APOE. Alzheimer's disease (AD) is characterized by progressive loss of memory and other cognitive abilities, as well brain atrophy and formation of amyloid plaques (Karch, Cruchaga, and Goate 2014). The APOE gene is involved in regulating lipid homeostasis. The e4 version of the APOE gene increases an individual's risk of late-onset Alzheimer disease (Liu et al. 2013).

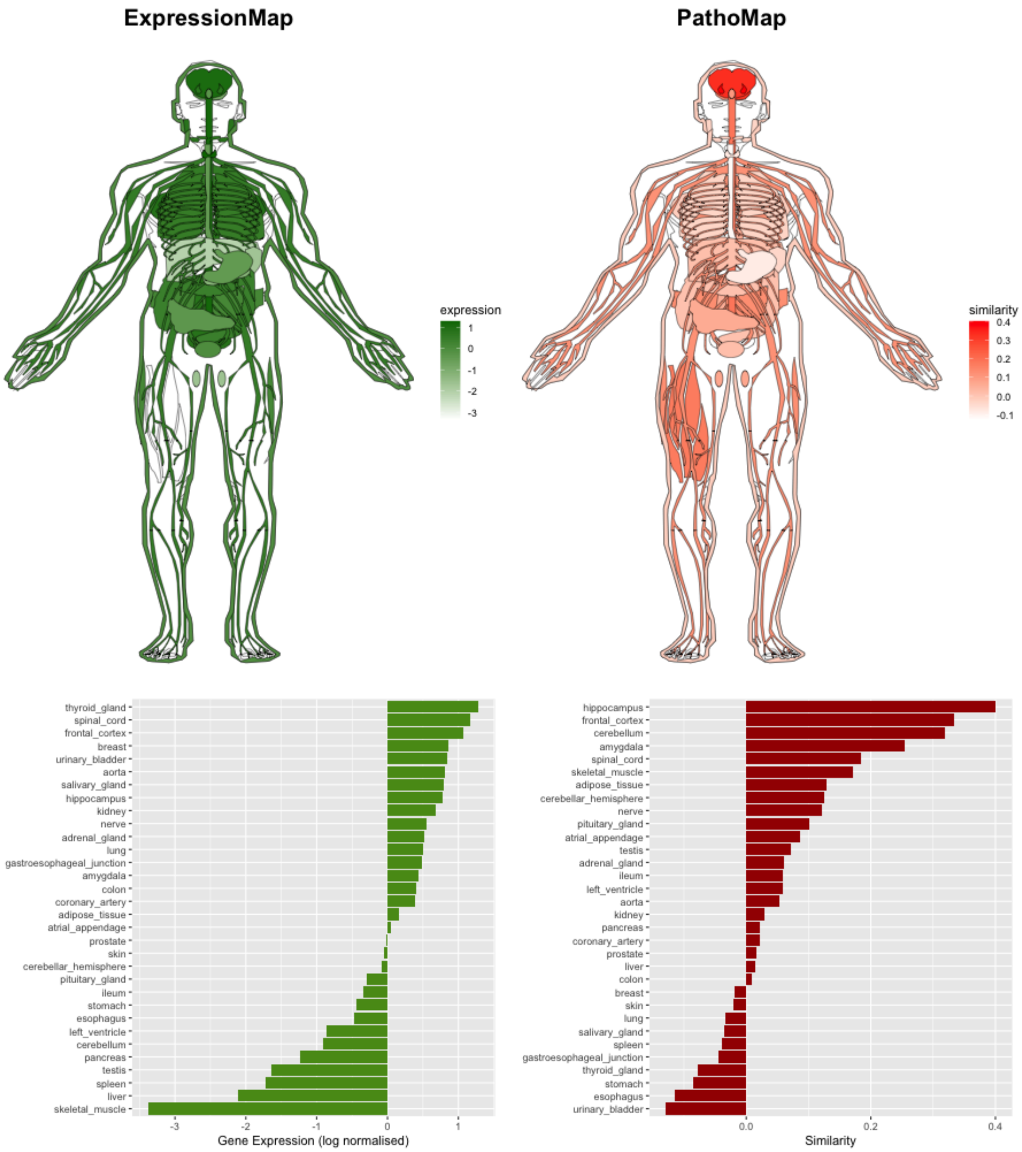


**Supplementary Figure S6.** Distribution of patho-scores and gene expression across organs for APP**.** Alzheimer's is a leading cause of dementia worldwide, and is characterized by production and accumulation of β-amyloid peptide (Aβ). The production of this neurotoxic peptide from sequential APP proteolysis is critical for progression of Alzheimer (O'Brien and Wong 2011) (Murphy and LeVine 2010). Early onset Alzheimer's disease is characterized by the presence of inherent dominant mutations in the APP gene (Karch, Cruchaga, and Goate 2014).

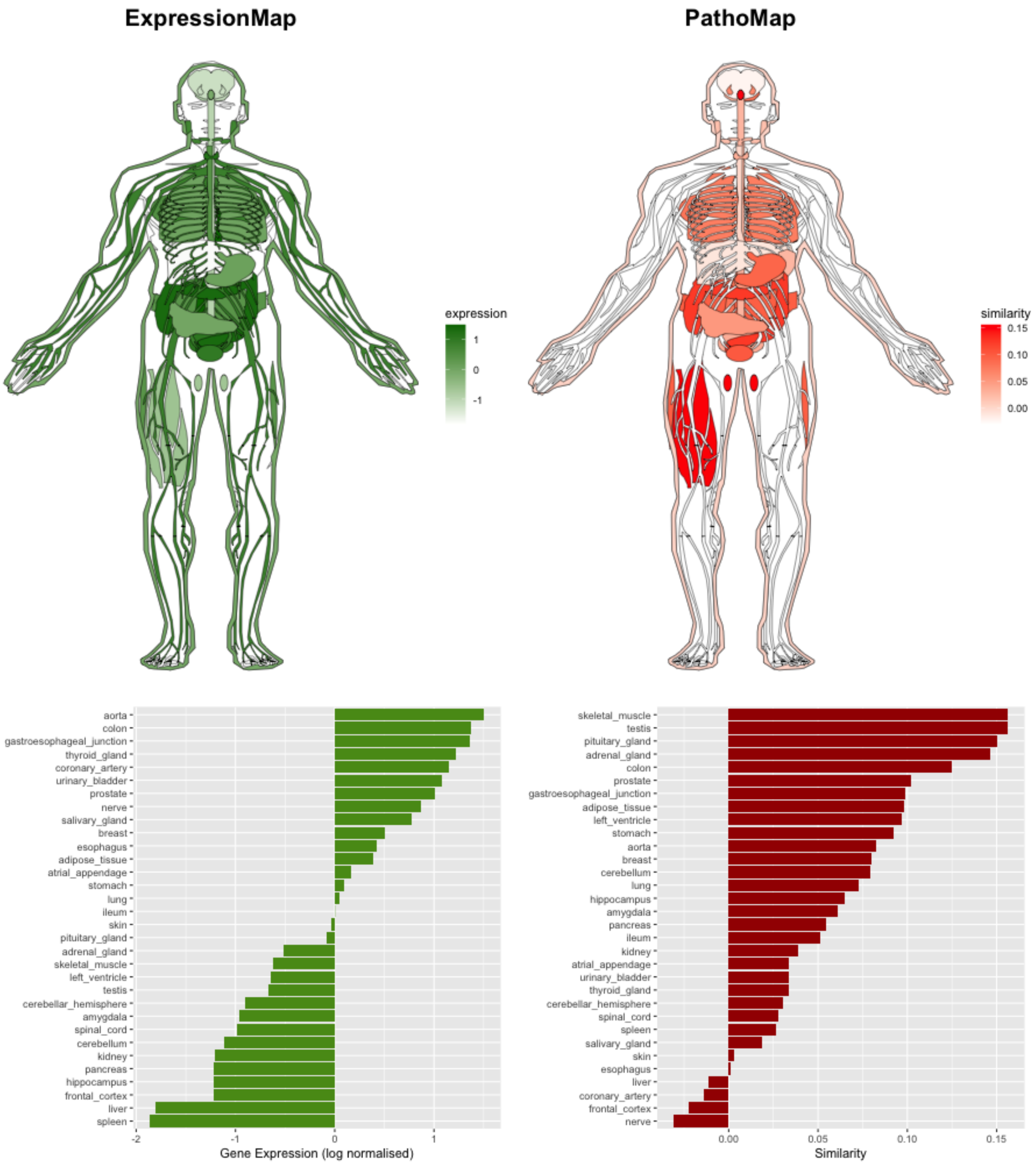


**Supplementary Figure S7.** Distribution of patho-scores and gene expression across organs for BMPR1A. Juvenile polyposis syndrome (JPS) is a rare autosomal dominant characterized by presence of multiple polyps in the gastrointestinal (GI) tract which increases risk of colon cancer. The germline mutations in BMPR1A causes JPS.
(Blatter et al. 2020) (Cichy, Klincewicz, and Plawski 2014).

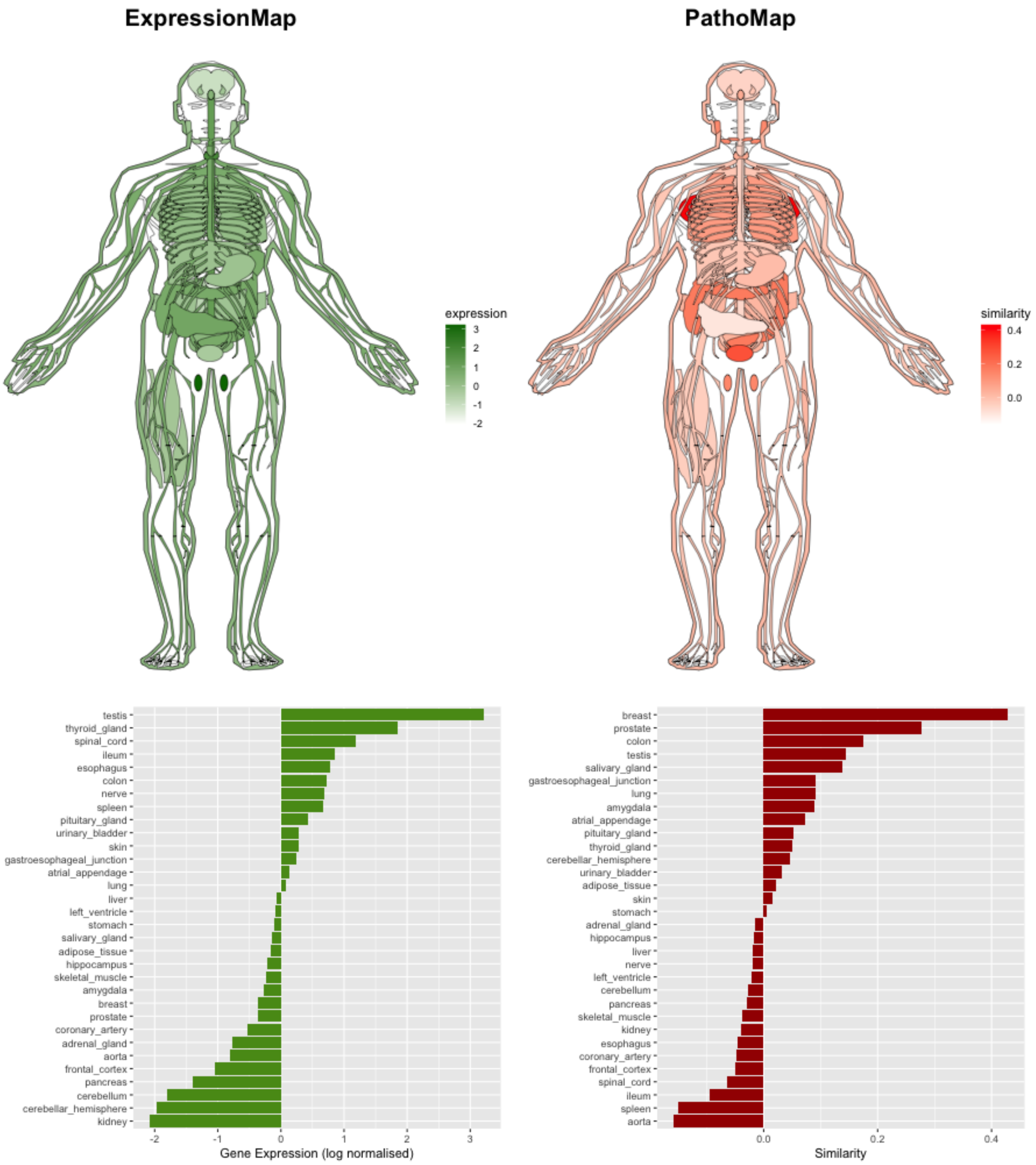


**Supplementary Figure S8.** Distribution of patho-scores and gene expression across organs for BRCA1**.** BRCA1 is a tumor suppressor gene. However, alterations or mutations in this gene predisposes individuals to increased risk of certain cancers such as breast, prostate cancer and other cancer types. (Godet and Gilkes 2017) (Levine et al. 2003).

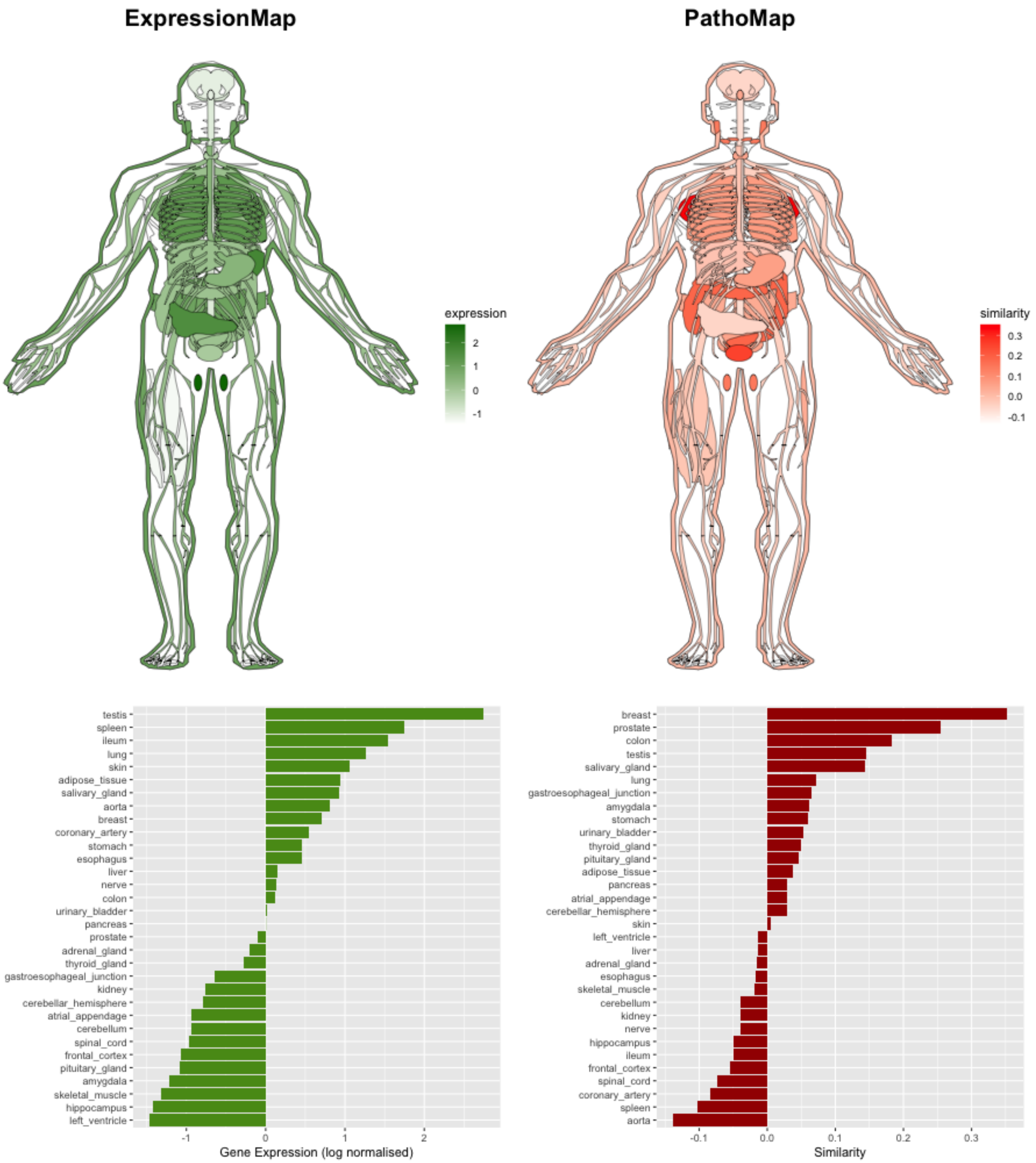


**Supplementary Figure S9.** Distribution of patho-scores and gene expression across organs for BRCA2. BRCA2 is a tumor suppressor gene. However, alterations or mutations in this gene predisposes individuals to increased risk of certain cancers such as breast, prostate cancer and other cancer types. (Godet and Gilkes 2017) (Levine et al. 2003).

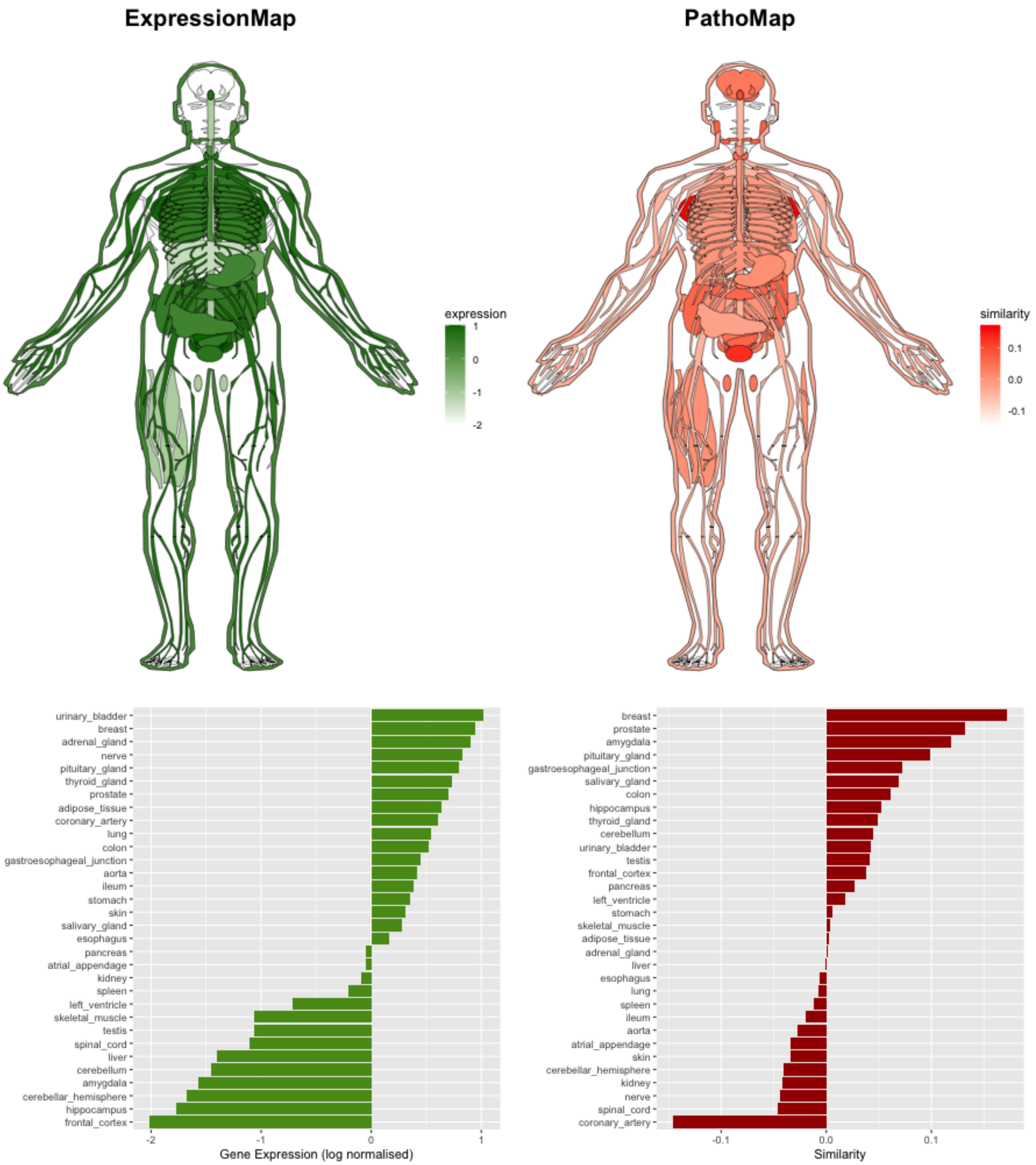


**Supplementary Figure S10.** Distribution of patho-scores and gene expression across organs for CDK4. Cyclin-dependent kinases (CDKs) regulate cell cycle checkpoint and transcriptional events, thus are key regulators of cell proliferation. Notably, dysregulation in CDKs results in uncontrolled cell proliferation. The CDK4 kinase is involved in initiation and maintenance of tumorigenesis in breast cancer (Ding et al. 2020).

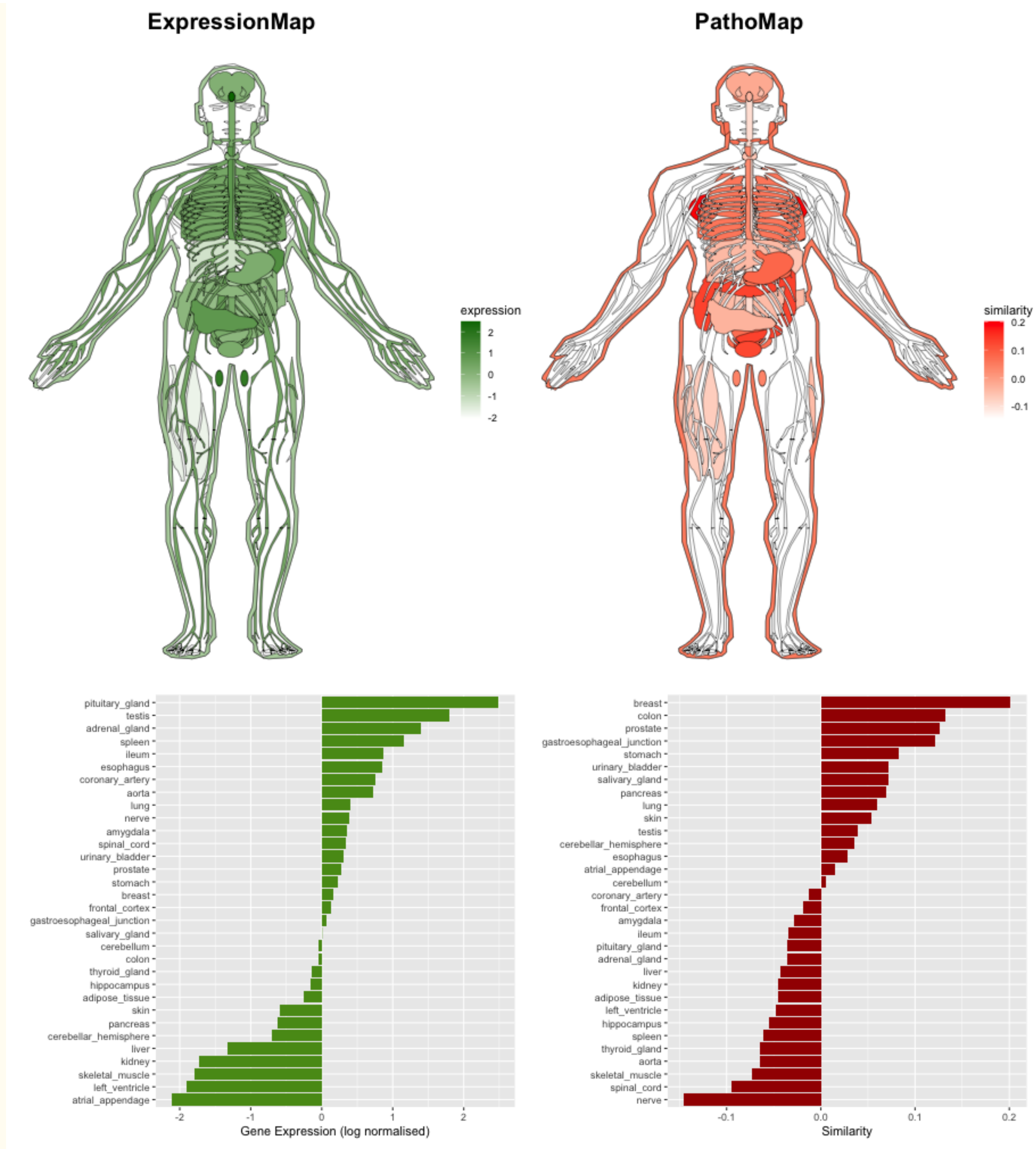


**Supplementary Figure S11.** Distribution of patho-scores and gene expression across organs for CDKN2A**.** CDKN2A is a tumor suppressor gene which encodes for p16INK4A and p14ARF proteins. CDKN2A gene is commonly inactivated due to genetic and epigenetic changes in multiple cancer types (Chan, Chiang, and Ngeow 2021). The mutations in the CDKN2A gene are associated with increased risk of pancreatic, melanoma and breast cancer (McWilliams et al. 2011).

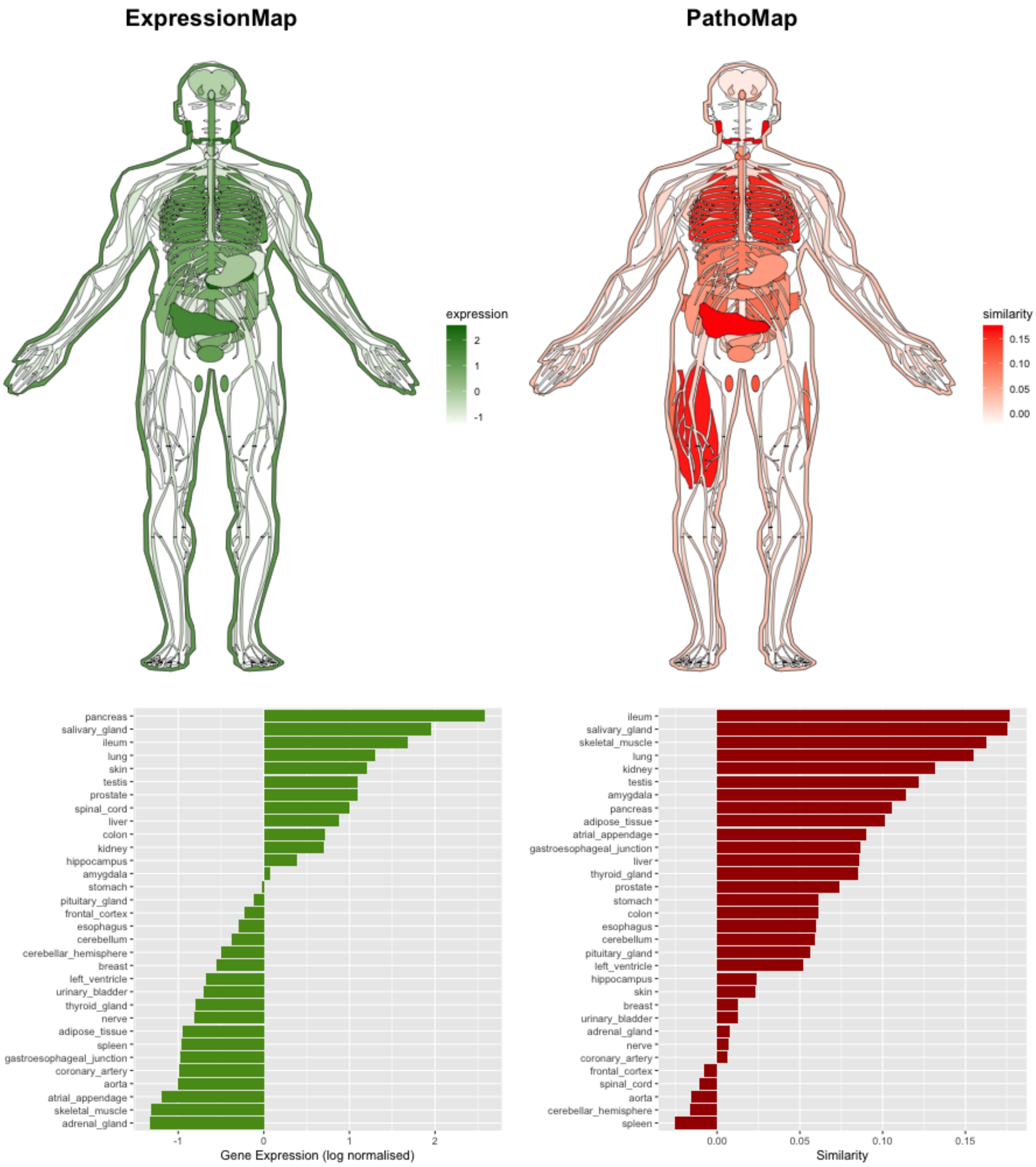


**Supplementary Figure S12.** Distribution of patho-scores and gene expression across organs for CFTR**.** Cystic fibrosis (CF) is an inherited lethal disorder caused by mutations in the CFTR gene (Mall and Hartl 2014). CFTR gene encodes for ion channels, dysfunction or alteration in this gene results in imbalance of ions and fluids in organs such as intestine, pancreas etc. (Lopes-Pacheco 2019).

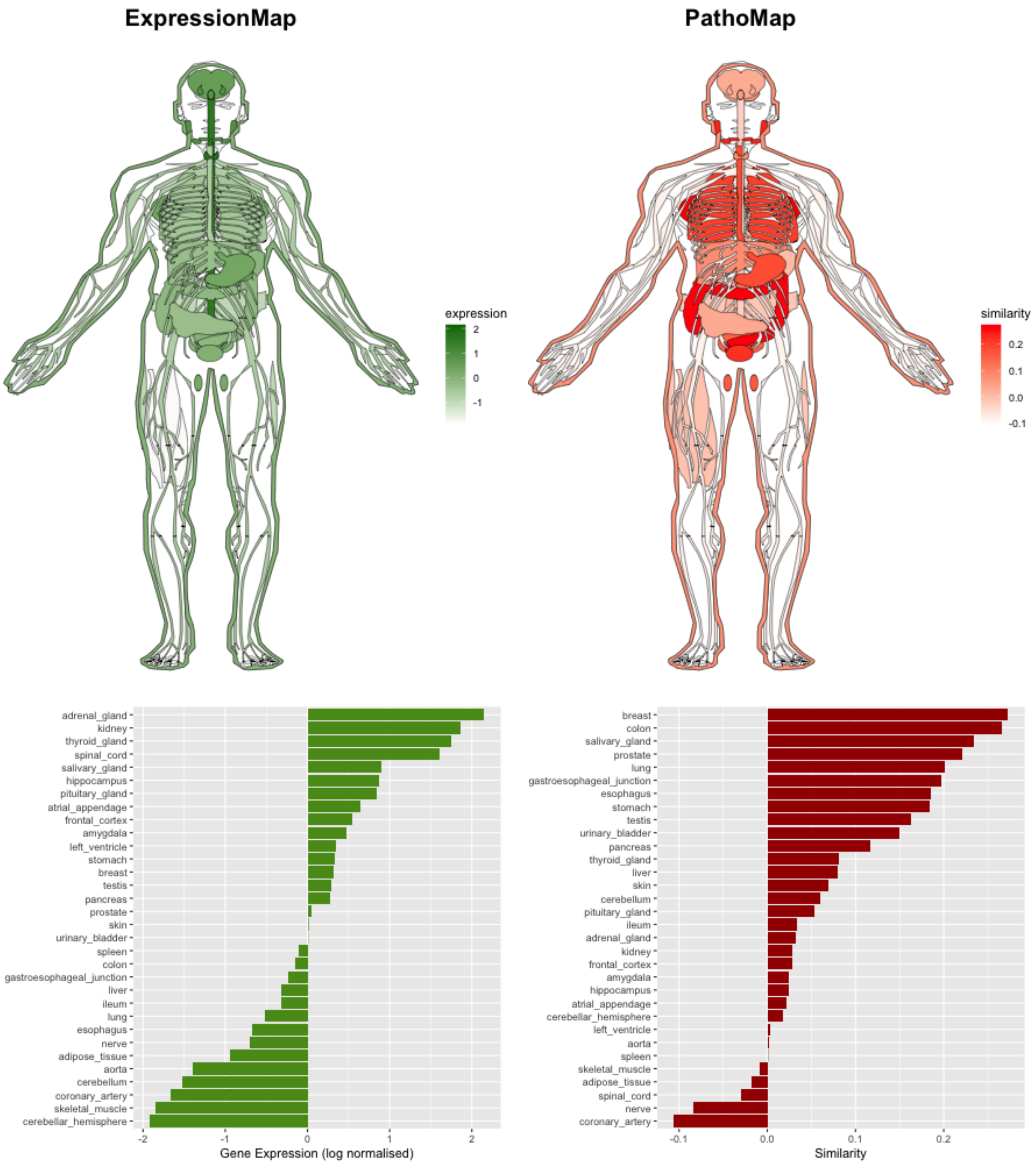


**Supplementary Figure S13.** Distribution of patho-scores and gene expression across organs for FHIT. FHIT is a potential tumor suppressor gene that is deleted or silenced in more than 50% of common cancer types (Kiss et al. 2017). In breast cancer, alterations in the FHIT gene are linked with genome instability (Ingvarsson 2001). The alterations in this gene have been reported in myriad of cancer types such a colon, esophageal, gastric and lung cancer (Silveira Zavalhia et al. 2018).

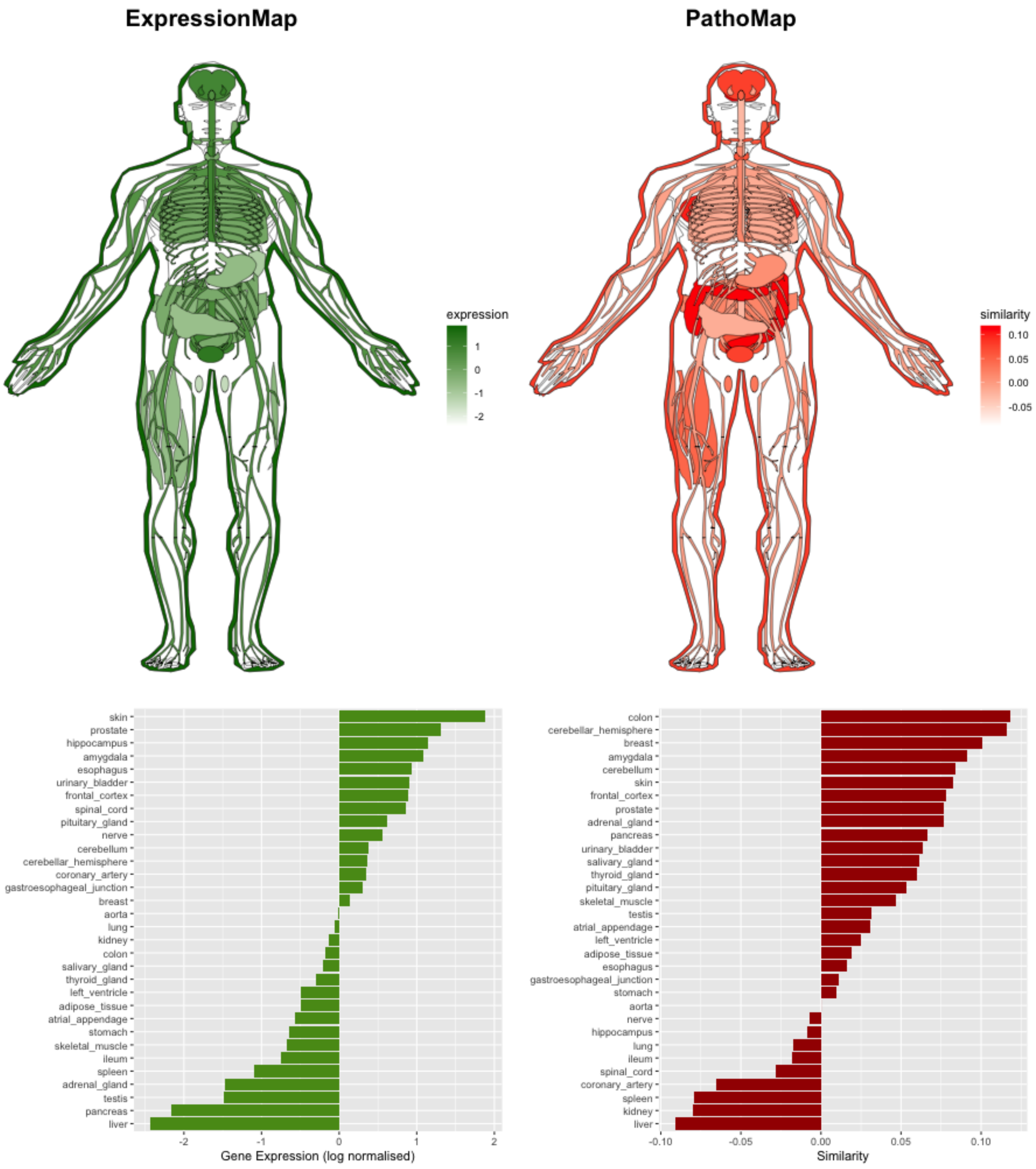


**Supplementary Figure S14.** Distribution of patho-scores and gene expression across organs for HRAS. HRAS is associated with diseases such as Costello Syndrome (Estep et al. 2006) and Epidermal Nevus Syndrome which involves skin related defects (Avitan-Hersh et al. 2014).

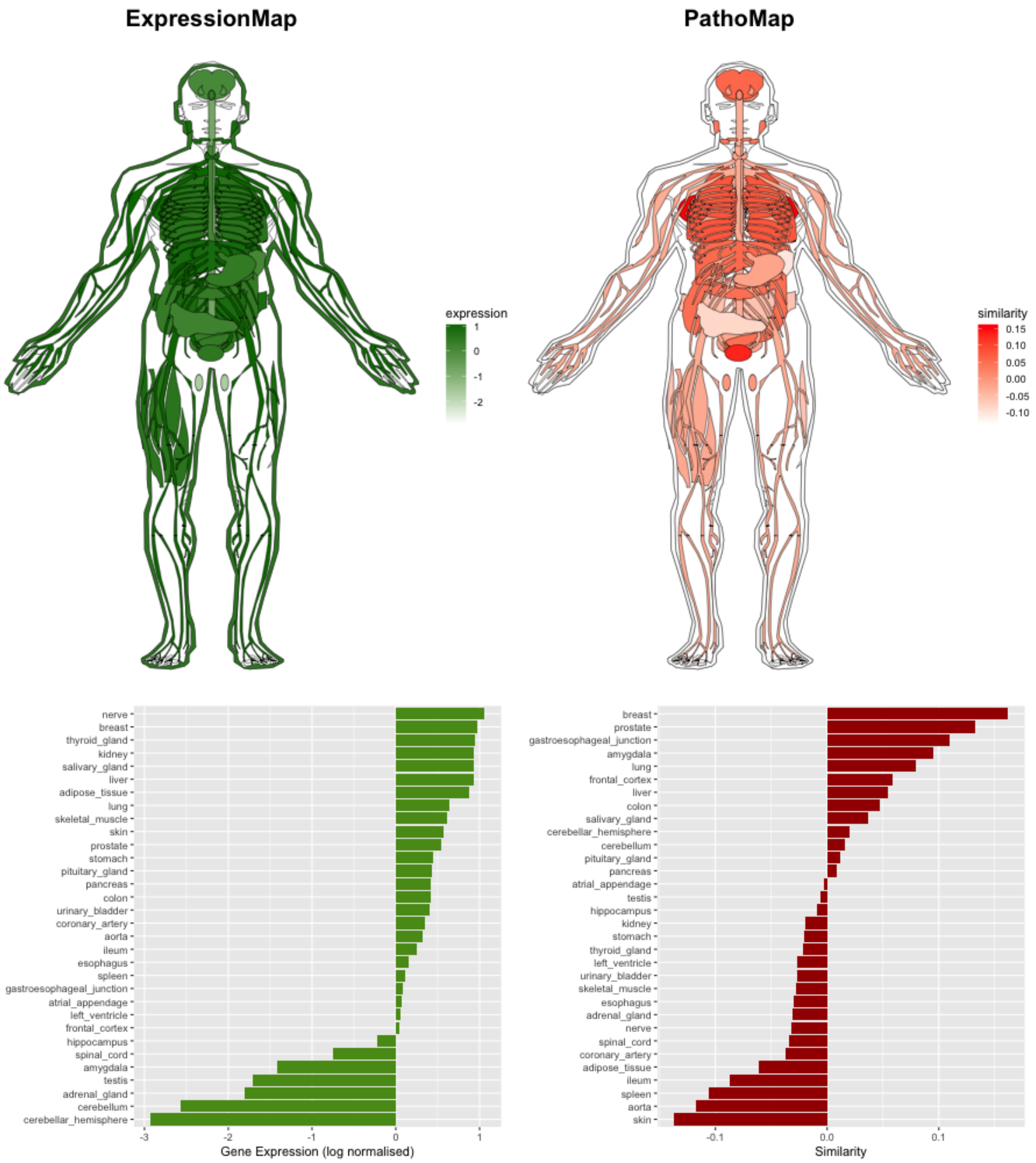


**Supplementary Figure S15.** Distribution of patho-scores and gene expression across organs for MET**.** Mesenchymal Epithelial Transition (MET) is a receptor tyrosine kinase activated by its ligand Hepatocyte Growth Factor (HGF). In many solid cancers, MET is mutated or over amplified. In prostate cancer, the expression of MET increases throughout the progression of prostate cancer and metastasis (Varkaris et al. 2011). Overexpression of MET is associated with poor survival outcomes in breast cancer patients (de Melo Gagliato et al. 2014).

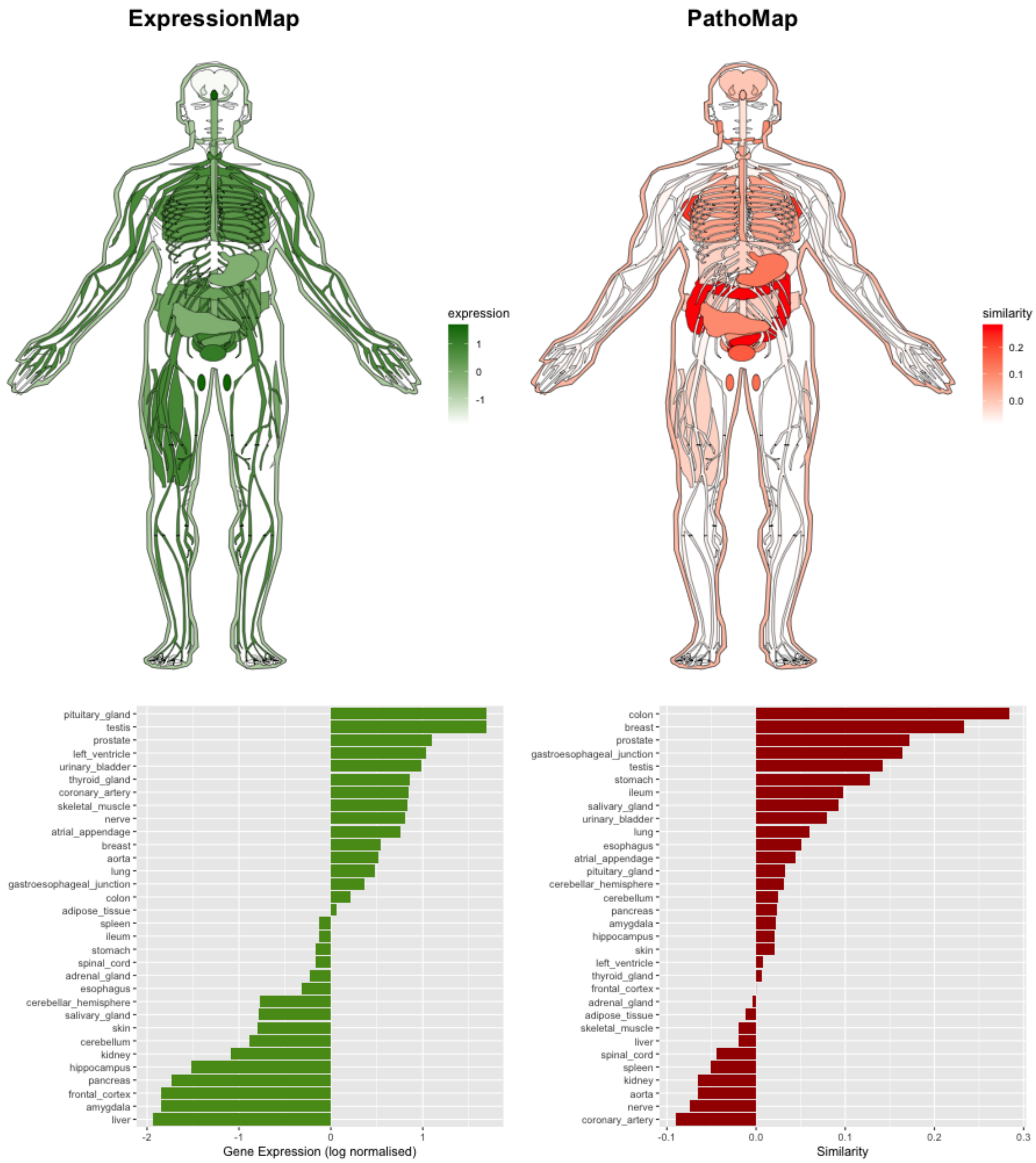


**Supplementary Figure S16.** Distribution of patho-scores and gene expression across organs for MLH1**.** MLH1 is a tumor suppressor gene known to play a role in DNA mismatch repair. Germline mutations in this gene are known to cause Lynch syndrome (LS). The germline mutations in this gene is associated with increased risk of colon, stomach and other known cancers (Win, Lindor, and Jenkins 2013). In hereditary breast cancer, frequency of gemline LS mutations including MLH1 is relatively high (Nikitin et al. 2020).

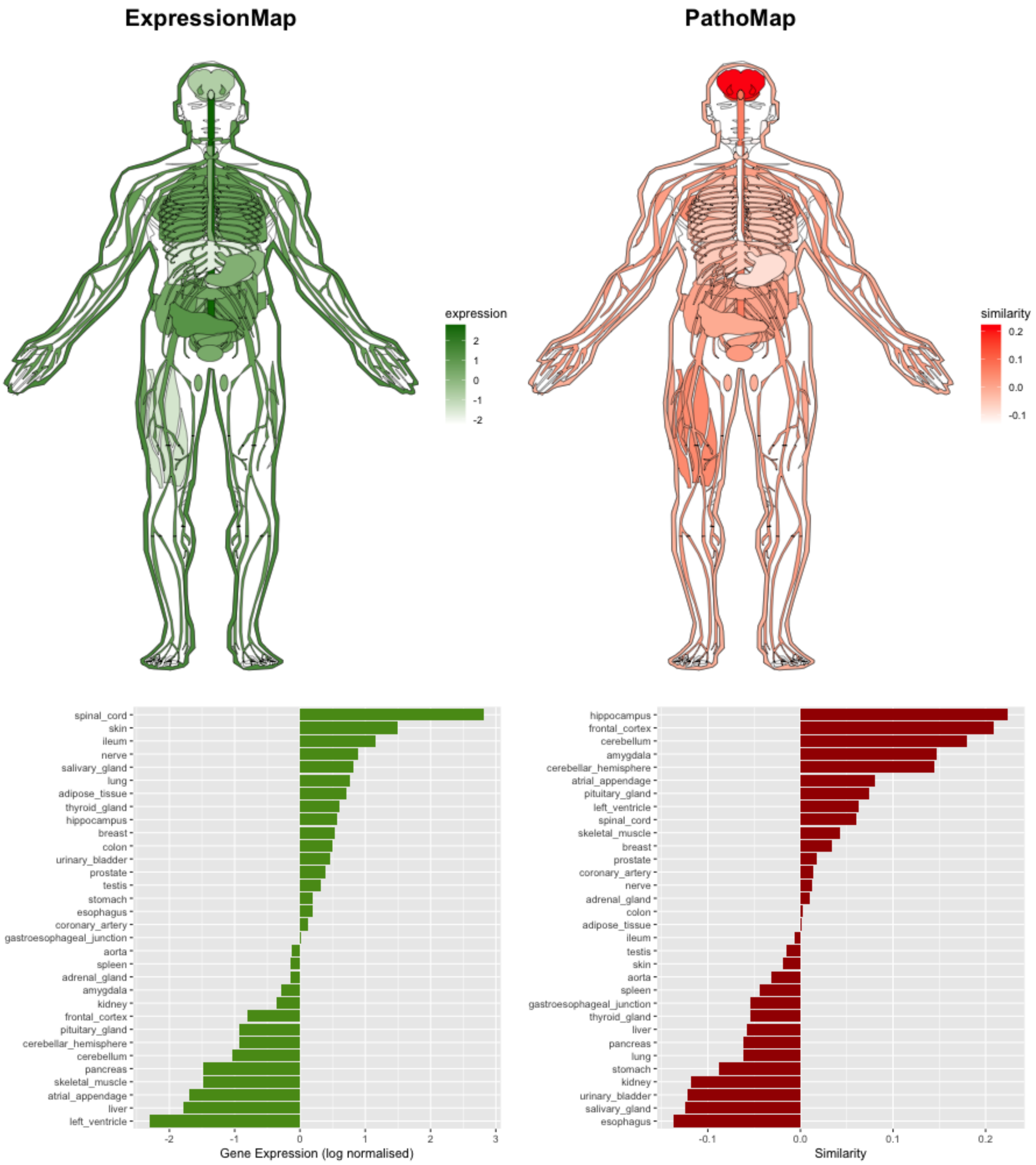


**Supplementary Figure S17.** Distribution of patho-scores and gene expression across organs for PSEN1**.** Mutations in the presenilin-1 (*PSEN1*) gene are involved in causing familial Alzheimer's disease and early-onset Alzheimer disease (Lanoiselée et al. 2017).

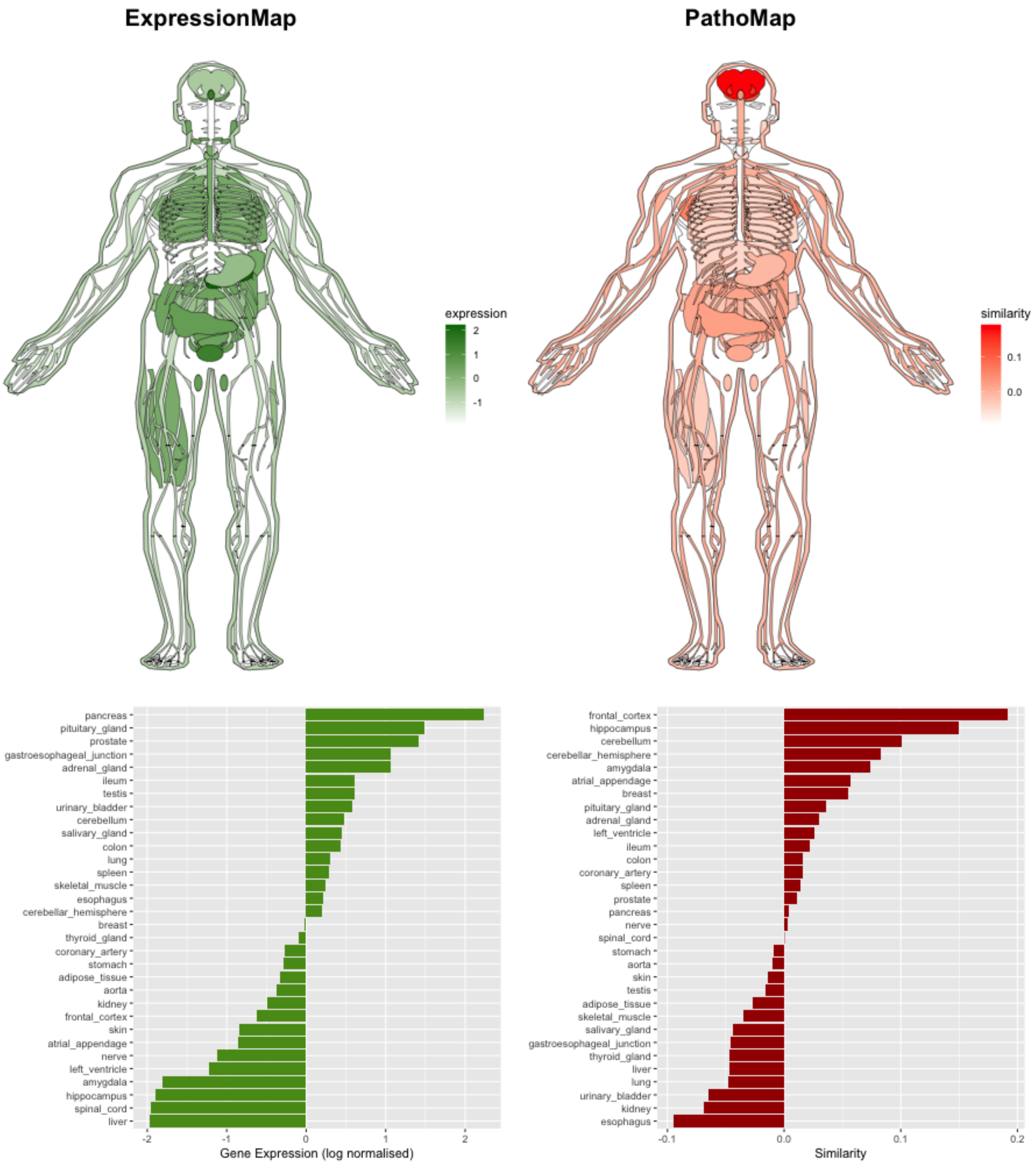


**Supplementary Figure S18.** Distribution of patho-scores and gene expression across organs for PSEN2**.** Mutations in the presenilin-2 (*PSEN2*) gene are known causes of familial Alzheimer's disease and are associated with early-onset Alzheimer disease (Lanoiselée et al. 2017) (An, Cai, and Kim 2015).

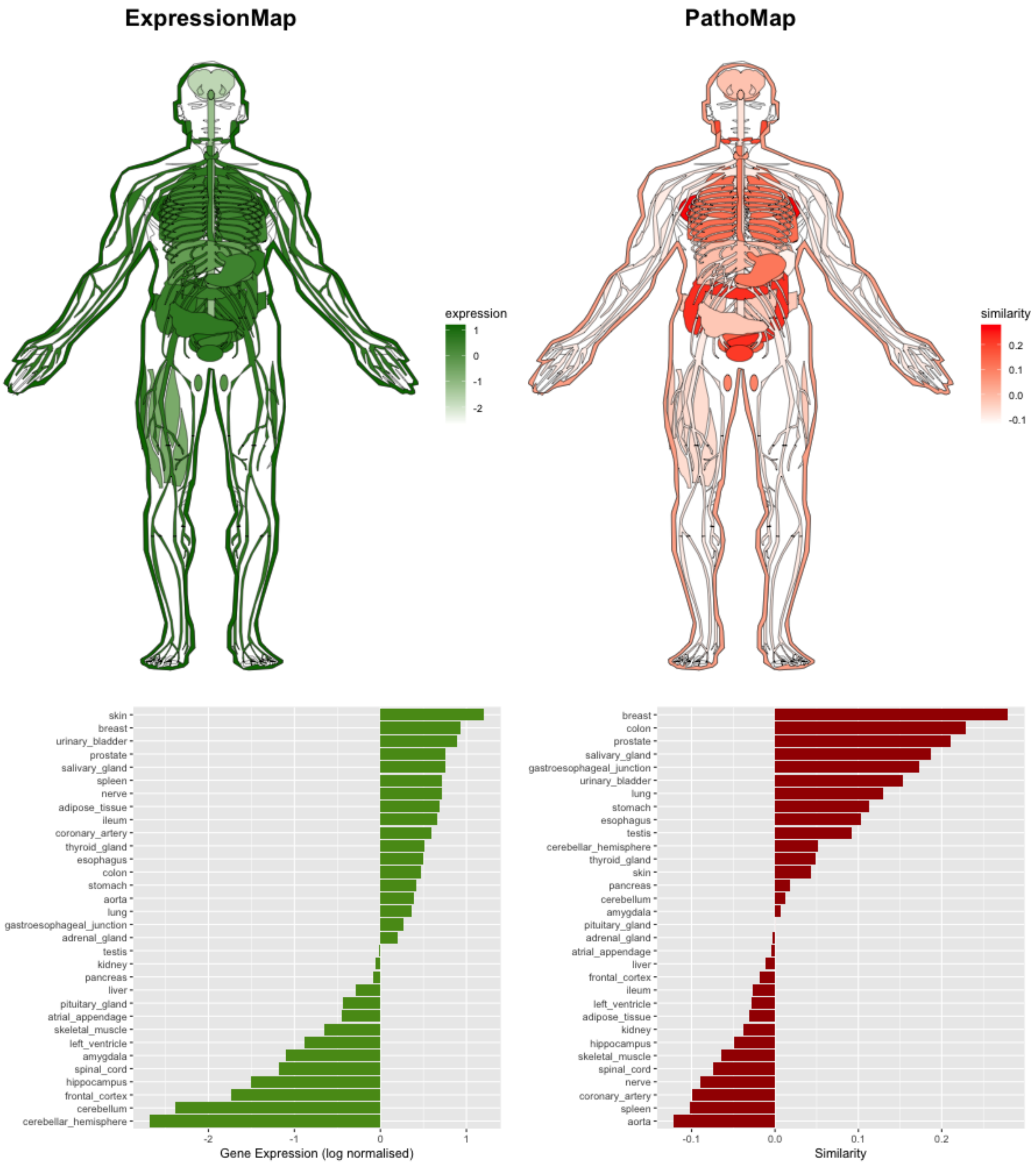


**Supplementary Figure S19.** Distribution of patho-scores and gene expression across organs for TP53**.** TP53 is a tumor suppressor gene which is found to be mutated in a number of cancers including breast cancer. A somatic mutation in the TP53 gene has been found in approximately 30% of breast tumours (Silwal-Pandit et al. 2014). TP53 alterations are also involved in driving development of colon cancer (Williams et al. 2020).

### *References*


An, Seong Soo, Yan Cai, and Sangyun Kim. 2015. "Mutations in Presenilin 2 and Its Implications in Alzheimer's Disease and Other Dementia-Associated Disorders." *Clinical Interventions in Aging*. https://doi.org/10.2147/cia.s85808.

Avitan-Hersh, Emily, Sameh Tatur, Margarita Indelman, Vardit Gepstein, Roni Shreter, Dov Hershkovitz, Riva Brick, Reuven Bergman, and Dov Tiosano. 2014. "Postzygotic HRAS Mutation Causing Both Keratinocytic Epidermal Nevus and Thymoma and Associated with Bone Dysplasia and Hypophosphatemia due to Elevated FGF23." *The Journal of Clinical Endocrinology and Metabolism* 99 (1): E132–36.

Blatter, Robert, Benjamin Tschupp, Stefan Aretz, Inge Bernstein, Chrystelle Colas, D. Gareth Evans, Maurizio Genuardi, et al. 2020. "Disease Expression in Juvenile Polyposis Syndrome: A Retrospective Survey on a Cohort of 221 European Patients and Comparison with a Literature-Derived Cohort of 473 SMAD4/BMPR1A Pathogenic Variant Carriers." *Genetics in Medicine: Official Journal of the American College of Medical Genetics* 22 (9): 1524–32.

Chan, Sock Hoai, Jianbang Chiang, and Joanne Ngeow. 2021. "CDKN2A Germline Alterations and the Relevance of Genotype-Phenotype Associations in Cancer Predisposition." *Hereditary Cancer in Clinical Practice* 19 (1): 21.

Cichy, Wojciech, Beata Klincewicz, and Andrzej Plawski. 2014. "Juvenile Polyposis Syndrome." *Archives of Medical Science: AMS* 10 (3): 570–77.

Ding, Lei, Jiaqi Cao, Wen Lin, Hongjian Chen, Xianhui Xiong, Hongshun Ao, Min Yu, Jie Lin, and Qinghua Cui. 2020. "The Roles of Cyclin-Dependent Kinases in Cell-Cycle Progression and Therapeutic Strategies in Human Breast Cancer." *International Journal of Molecular Sciences* 21 (6). https://doi.org/10.3390/ijms21061960.

Estep, Anne L., William E. Tidyman, Michael A. Teitell, Philip D. Cotter, and Katherine A. Rauen. 2006. "HRAS Mutations in Costello Syndrome: Detection of Constitutional Activating Mutations in Codon 12 and 13 and Loss of Wild-Type Allele in Malignancy." *American Journal of Medical Genetics. Part A* 140 (1): 8–16.

Fodde, R. 2002. "The APC Gene in Colorectal Cancer." *European Journal of Cancer* 38 (7): 867–71.

Godet, Inês, and Daniele M. Gilkes. 2017. "BRCA1 and BRCA2 Mutations and Treatment Strategies for Breast Cancer." *Integrative Cancer Science and Therapeutics* 4 (1). https://doi.org/10.15761/ICST.1000228.

Ingvarsson, S. 2001. "FHIT Alterations in Breast Cancer." *Seminars in Cancer Biology* 11 (5): 361–66.

Karch, Celeste M., Carlos Cruchaga, and Alison M. Goate. 2014. "Alzheimer's Disease Genetics: From the Bench to the Clinic." *Neuron*. https://doi.org/10.1016/j.neuron.2014.05.041.

Kiss, Daniel L., William Baez, Kay Huebner, Ralf Bundschuh, and Daniel R. Schoenberg. 2017. "Impact of FHIT Loss on the Translation of Cancer-Associated mRNAs." *Molecular Cancer* 16 (1): 179.

Lanoiselée, Hélène-Marie, Gaël Nicolas, David Wallon, Anne Rovelet-Lecrux, Morgane Lacour, Stéphane Rousseau, Anne-Claire Richard, et al. 2017. "APP, PSEN1, and PSEN2 Mutations in Early-Onset Alzheimer Disease: A Genetic Screening Study of Familial and Sporadic

Cases." *PLoS Medicine* 14 (3): e1002270.
Leoz, Maria Liz, Sabela Carballal, Leticia Moreira, Teresa Ocaña, and Francesc Balaguer. 2015. "The Genetic Basis of Familial Adenomatous Polyposis and Its Implications for Clinical Practice and Risk Management." *The Application of Clinical Genetics* 8 (April): 95–107.
Levine, Douglas A., Peter A. Argenta, Cindy J. Yee, David S. Marshall, Narciso Olvera, Faina Bogomolniy, Jamal A. Rahaman, et al. 2003. "Fallopian Tube and Primary Peritoneal Carcinomas Associated with BRCA Mutations." *Journal of Clinical Oncology: Official Journal of the American Society of Clinical Oncology* 21 (22): 4222–27.
Liu, Chia-Chen, Chia-Chan Liu, Takahisa Kanekiyo, Huaxi Xu, and Guojun Bu. 2013. "Apolipoprotein E and Alzheimer Disease: Risk, Mechanisms and Therapy." *Nature Reviews. Neurology* 9 (2): 106–18.
Lopes-Pacheco, Miquéias. 2019. "CFTR Modulators: The Changing Face of Cystic Fibrosis in the Era of Precision Medicine." *Frontiers in Pharmacology* 10: 1662.
Mall, Marcus A., and Dominik Hartl. 2014. "CFTR: Cystic Fibrosis and beyond." *The European Respiratory Journal: Official Journal of the European Society for Clinical Respiratory Physiology* 44 (4): 1042–54.
McWilliams, Robert R., Eric D. Wieben, Kari G. Rabe, Katrina S. Pedersen, Yanhong Wu, Hugues Sicotte, and Gloria M. Petersen. 2011. "Prevalence of CDKN2A Mutations in Pancreatic Cancer Patients: Implications for Genetic Counseling." *European Journal of Human Genetics: EJHG* 19 (4): 472–78.
Melo Gagliato, Debora de, Denis L. Fontes Jardim, Gerald Falchook, Chad Tang, Ralph Zinner, Jennifer J. Wheler, Filip Janku, et al. 2014. "Analysis of MET Genetic Aberrations in Patients with Breast Cancer at MD Anderson Phase I Unit." *Clinical Breast Cancer* 14 (6): 468–74.
Murphy, M. Paul, and Harry LeVine 3rd. 2010. "Alzheimer's Disease and the Amyloid-Beta Peptide." *Journal of Alzheimer's Disease: JAD* 19 (1): 311–23.
Nikitin, Aleksey G., Daria A. Chudakova, Rafael F. Enikeev, Dina Sakaeva, Maxim Druzhkov, Leyla H. Shigapova, Olga I. Brovkina, Elena I. Shagimardanova, Oleg A. Gusev, and Marat G. Gordiev. 2020. "Lynch Syndrome Germline Mutations in Breast Cancer: Next Generation Sequencing Case-Control Study of 1,263 Participants." *Frontiers in Oncology* 10 (May): 666.
O'Brien, Richard J., and Philip C. Wong. 2011. "Amyloid Precursor Protein Processing and Alzheimer's Disease." *Annual Review of Neuroscience* 34: 185–204.
Silveira Zavalhia, Lisiane, Aline Weber Medeiros, Andrew Oliveira Silva, and Adriana Vial Roehe. 2018. "Do FHIT Gene Alterations Play a Role in Human Solid Tumors?" *Asia-Pacific Journal of Clinical Oncology* 14 (5): e214–23.
Silwal-Pandit, Laxmi, Hans Kristian Moen Vollan, Suet-Feung Chin, Oscar M. Rueda, Steven McKinney, Tomo Osako, David A. Quigley, et al. 2014. "TP53 Mutation Spectrum in Breast Cancer Is Subtype Specific and Has Distinct Prognostic Relevance." *Clinical Cancer Research: An Official Journal of the American Association for Cancer Research* 20 (13): 3569–80.
Varkaris, Andreas, Paul G. Corn, Sanchaika Gaur, Farshid Dayyani, Christopher J. Logothetis, and Gary E. Gallick. 2011. "The Role of HGF/c-Met Signaling in Prostate Cancer Progression and c-Met Inhibitors in Clinical Trials." *Expert Opinion on Investigational Drugs* 20 (12): 1677–84.

Williams, David S., Dmitri Mouradov, Clare Browne, Michelle Palmieri, Meg J. Elliott, Rebecca Nightingale, Catherine G. Fang, et al. 2020. “Overexpression of TP53 Protein Is Associated with the Lack of Adjuvant Chemotherapy Benefit in Patients with Stage III Colorectal Cancer.” *Modern Pathology: An Official Journal of the United States and Canadian Academy of Pathology, Inc* 33 (3): 483–95.

Win, Aung Ko, Noralane M. Lindor, and Mark A. Jenkins. 2013. “Risk of Breast Cancer in Lynch Syndrome: A Systematic Review.” *Breast Cancer Research: BCR* 15 (2): R27.
